# Feature interference as a neuronal basis for the behavioural cost of task uncertainty

**DOI:** 10.1101/2024.03.04.583375

**Authors:** Cheng Xue, Sol K. Markman, Ruoyi Chen, Lily E. Kramer, Marlene R. Cohen

## Abstract

Humans and animals have an impressive ability to juggle multiple tasks in a constantly changing environment. This flexibility, however, worsens performance under uncertain task conditions. Here, we combined monkey electrophysiology, human psychophysics, and artificial neural network modeling to investigate the neuronal mechanisms underlying this performance cost. We developed a behavioural paradigm to measure and influence participants’ decision-making and perception in two distinct perceptual tasks. Our data reveal that both humans and monkeys, unlike an artificial neural network trained for the same tasks, make less accurate perceptual decisions when the task is uncertain. We generated a mechanistic hypothesis by comparing this neural network trained to produce correct choices with another network trained to replicate the participants’ choices. We hypothesized, and confirmed with further behavioural, physiological, and causal experiments, that the cost of task uncertainty and flexibility comes from what we term feature interference. Under uncertain conditions, interference between different tasks causes errors because it results in stronger representations of irrelevant features and entangled neuronal representations of different features. Our results suggest a tantalizing, general hypothesis: that cognitive capacity limitations, both in health and disease, stem from interference between neural representations of different stimuli, tasks, or memories.

## Introduction

In the face of the inherent uncertainty and unexpected changes in natural environments, humans and animals have evolved remarkable cognitive flexibility, enabling them to seamlessly switch between tasks as the situation demands. Such flexibility is an integral part of natural decision-making (*1*), but comes with a cost: when we are uncertain which task should be performed (typically during and shortly before or after switching to a different task), we take longer to perform the task we choose and do so with lower accuracy (*1–4*).

Numerous non-exclusive hypotheses have been proposed based on behavioural evidence to explain the worse and slower choices during task switching. These include limits on processing capacity (*4*), undue attention to irrelevant information (*5–8*), inhibition of relevant information (*2*), task-set reconfiguration (*2*), and proactive interference between competing tasks (*2, 3, 9–12*). These phenomenological explanations of behaviour may reflect a common neuronal mechanism leading to poorer perceptual decisions and longer reaction times during task switching.

Here, we propose a novel neural mechanism that integrates and elucidates the roles of various hypothesized reasons for the cost of task uncertainty and flexibility. We used a multidisciplinary approach including behavioural experiments in humans and rhesus monkeys, multi-neuron, multi-area recordings and causal manipulations in monkeys, and artificial neural network modeling to generate and test neural hypotheses. The core of our approach is our two-feature visual discrimination task, in which participants must discriminate one visual feature and ignore the other, inferring the implicitly changing task rule that determines which feature is relevant, based on their history of stimuli, choices and rewards. By measuring and manipulating task certainty, we can directly link the corresponding behavioural adaptations to neural computations in both behaving animals and artificial networks, providing more comprehensive insights into the neuronal basis of cognitive flexibility.

Our findings suggest that the cost of task flexibility arises from stronger interference between representations of multiple features in contexts where the task-relevance of the features is not certain. Such interference is a joint product of retaining task-irrelevant visual information and the entanglement of the neural representations of unrelated features. We hypothesize that this phenomenon underlies the cost of cognitive flexibility across species, systems, and health states, offering promising avenues for advancing our understanding and addressing cognitive limitations in both basic science and translational research.

## Results

### Worse perceptual performance under task uncertainty

We designed a two-feature discrimination task that allows us to measure and manipulate perceptual difficulty and task certainty. In this paradigm, participants perform the same perceptual discriminations while task certainty (certainty about which task to perform) varies based on the stimulus, choice, and reward history (*1*) (Fig. 1A). We trained 206 online human participants and two rhesus monkeys (Macaca mulatta; both male, 12 and 9 kg) to discriminate changes in two visual features of subsequently presented Gabor stimuli (spatial frequency and orientation for human participants; spatial frequency and location for monkey participants). On each trial, both features changed at magnitudes near perceptual threshold (average perceptual performance is 79% for monkeys, and 82% for humans), but only one was relevant. Participants indicated both their chosen task and perceptual discrimination on every trial, and only correct discriminations of the relevant feature were rewarded (Fig. 1A-B). Because the reward outcome of a choice is completely independent of the feature deemed irrelevant, there is no benefit in maintaining the irrelevant feature’s representation, either for obtaining a reward on the current trial or for inferring the correct task. The separation of response targets for the two tasks allowed us direct access to the participants’ subjective judgement of the task rule on every trial, independent of the objective task rule enforced on every trial.

**Fig 1.**
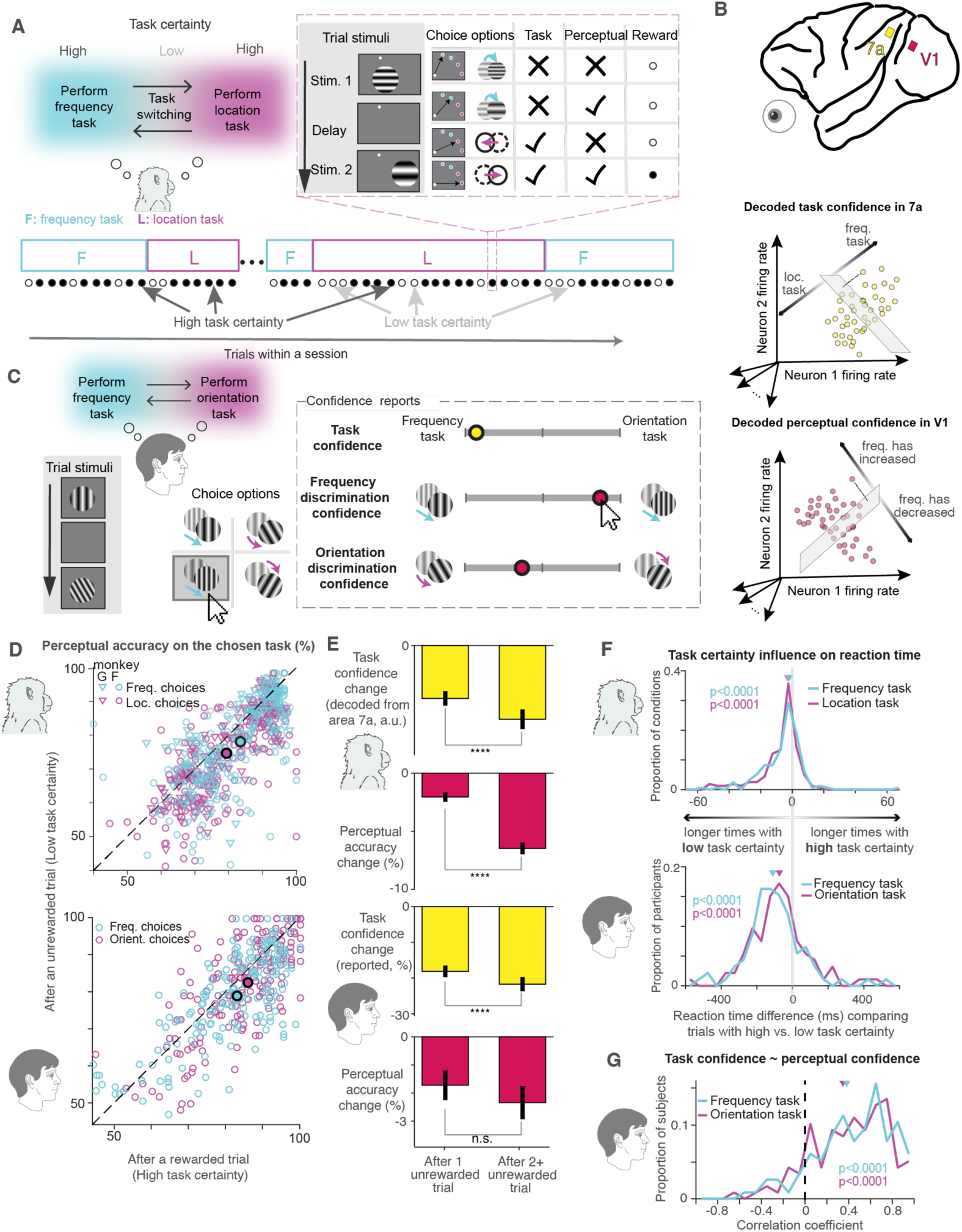
Tasks and the behavioural cost of task uncertainty. (**A**) Monkeys discriminated either the spatial frequency (cyan) or location (magenta) of two sequentially presented Gabor stimuli while ignoring changes in the irrelevant feature. The relevant feature is not explicitly cued. There was a fixed, low probability of a task switch after each trial, leading to task blocks of variable length (cyan and magenta rectangles). Participants had to infer the relevant feature from their history of choices and feedback. Participants were rewarded only if they made the correct task choice and perceptual judgment (they indicated both on each trial by making a saccade to one of four possible visual targets on screen that represent the choices schematized in the first column of the table in A). Therefore, task certainty was generally high following rewarded trials and low following unrewarded trials. (**B**) Electrophysiological recording in macaque cortical area V1 area 7a (top). Crimson and yellow squares represent approximate locations for the Utah arrays used for recording in each area. The monkeys’ task confidence was inferred from 7a activity: we identified a hyperplane discriminating chosen tasks in neural response space, taking the Euclidean distance from each trial’s activity to this hyperplane as confidence (middle). Similarly, perceptual confidence was inferred from V1 activity (bottom) (**C**) Human participants performed the same task as the monkeys, alternating between orientation discrimination and frequency discrimination. On a small proportion of trials, some human participants were asked to report their confidence about their task choice and perceptual discrimination of both features on a continuous scale. (**D**) Behavioural cost of flexibility in perceptual performance across sessions and participants for monkeys (top) and humans (bottom). Each open symbol represents perceptual performance for a set of trials with identical feature change amounts within a session. Both species show worse perceptual performance under low task certainty (following an unrewarded trial; y-axis) than under high task certainty (following a rewarded trial; x-axis; p=0.003 for human orientation choices, p<0.0001 for human frequency choices; p<0.0001 for monkey location and frequency choices. Filled symbols denote mean performance. (**E**) Task confidence and perceptual performance drop following unrewarded trials. The bar plots report changes in task confidence (yellow bars) and perceptual performance (crimson bars) following a single unrewarded trial (left bars) and following multiple consecutive unrewarded trials (right bars), all relative to the task confidence or perceptual performance after rewarded trials, so negative values indicate weaker task confidence / poorer perceptual performance after unrewarded trials compared to those after rewarded trials. Error bars indicate s.e.m. Monkeys’ task confidence, decoded from parietal area 7a neuronal activity, decreased significantly after unrewarded trials (upper yellow bars, p<0.0001 for both), with significantly more decrease following consecutive unrewarded trials than that following a single unrewarded trial (p<0.0001). Similarly, human participants’ task confidence reports showed the same pattern (lower yellow bars, both significantly below 0, p<0.0001, and significantly different with each other, p<0.0001). Perceptual performance in both species decreased with task uncertainty (all crimson bars show distributions significantly below 0, monkeys p=0.0002 after single unrewarded trial, p<0.0001 after multiple consecutive unrewarded trials; humans p=0.03 after single unrewarded trial, p=0.00018 after multiple unrewarded trials). Perceptual performance declined even more after multiple consecutive unrewarded trials than after a single non-reward for monkeys (p<0.0001) but not for humans (p=0.13). (**F**) Cost of task uncertainty on reaction time. The upper (monkey) and lower (human) histograms show distributions of the differences between reaction times under high and low task certainty for matched stimuli. The histograms are shifted to the left, indicating that even when participants correctly discriminated the feature change, it took them longer under low task certainty than under high task certainty. p<0.0001 for both species and both tasks. (**G**) Behavioural cost of flexibility in confidence. Cyan and magenta curves show significantly positive distributions of correlation coefficients between task confidence and perceptual confidence on the chosen task for human participants (p<0.0001 for both tasks), which means that overall, when participants’ task confidence is low, their confidence on task performance is also low.

The participants need to infer the rule change based on recent stimulus, choice, and reward history. The relevant feature switched without a cue with a constant low probability on each trial (2.5% for monkeys and 10% for humans). A reward/positive feedback was given only when the participant both correctly identified the relevant feature to discriminate and performed that discrimination correctly. Therefore, on trials that follow a reward, participants could be fairly certain that the rewarded task would be the same as the previous trial (97.5% certain for monkeys, 90% certain for humans). In contrast, a non-reward is ambiguous: it could reflect an error in task selection, perceptual discrimination, or both. As a proxy for task certainty, we therefore compare behavior and neuronal responses on trials that follow rewarded or un-rewarded trials in many analyses.

While the monkeys performed this task, we simultaneously recorded populations of neurons in their primary visual cortex (area V1) and parietal cortex (area 7a) (Fig. 1B). On a subset of trials, human participants were asked to answer additional questions, such as perceptual and task confidence (Fig. 1C).

Our participants reacted to task switches in a close to normative fashion. Similar to an ideal observer, they inferred new task rules from missed rewards and restored reward rates within a few trials (Fig. S1A, left panel). Errors in which participants missed a task change occur at a much higher rate than the rate of “false alarms” (when participants incorrectly switch tasks despite no task change, which would be consistent with exploratory behavior). This mismatch indicates a conservative threshold for task switching, which is qualitatively consistent with ideal observer behaviour (Fig. S1B). These results suggest that the participants’ task switches largely did not reflect random exploration between the two tasks, but likely reflect the accumulation of evidence for a task change.

Consistent with an accumulation process, the participants’ task uncertainty increases with multiple consecutive non-rewards. Human participants’ confidence about the relevant task was lower after consecutive unrewarded trials than after a single unrewarded trial (Fig. 1E lower yellow bar plot) (Fig. 1E, third panel from the top). Similarly, inferring the monkeys’ task confidence from the representation of task choices in parietal area 7a suggests their task confidence followed a similar pattern: the neuronal inference of confidence was decreased more after consecutive unrewarded trials than after a single non-reward (Fig. 1E, first panel from the top).

Several additional lines of evidence indicate that our participants understand the structure of the task well enough to incorporate the strength of sensory evidence into their updating of task certainty. The monkeys were more likely to switch tasks after unrewarded trials with big, obvious changes in the deemed relevant feature, which offer stronger evidence that the task has changed (Fig. S1C). Even among unrewarded trials with exactly the same feature changes, task switches were more likely when V1 neurons indicated stronger perceptual confidence (Figure S1D). As a simple proxy for these many measures of task certainty, for most analyses we compare behaviour and neural responses on trials that were immediately preceded by a rewarded trial or an unrewarded trial.

Both humans and monkeys exhibit behavioural costs during task switching, reflected in both perceptual performance and response times. The reward rate and perceptual performance (i.e. whether they made the correct perceptual choice, regardless of whether the chosen task was correct) for monkey participants both drop following task-rule changes (Fig. S1A red lines). While the drop in reward rate following task-rule change is expected even for an ideal observer (Fig. S1A, left panel, blue line), the perceptual performance drop is not normative: since visual feature changes were drawn at random on each trial, the perceptual judgments should not depend on trial history (Fig. S1A, right panel, blue line). Yet, perceptual performances for monkeys and humans significantly declined following unrewarded trials (i.e. high task uncertainty) compared to rewarded trials (i.e. minimal task uncertainty) (Fig. 1D). In both species, perceptual performance decreased along with task confidence following both a single unrewarded trial or multiple unrewarded trials (Fig. 1E). In monkeys, perceptual performances worsened further after consecutive unrewarded trials (i.e. even higher task uncertainty) (Fig. 1E, upper crimson bar plot), which shows that higher levels of task uncertainty are associated with more decline in perceptual performance.

In addition to performance declines, both humans and monkeys responded more slowly under high task uncertainty (Fig. 1F, Fig. S1E). Both human and monkey participants show a behavioural cost of flexibility in perceptual confidence as well. In humans, task confidence is positively correlated with perceptual confidence (Fig. 1E). This is consistent with the finding that in monkeys, trial-to-trial correlations between task and perceptual confidence, decoded from area 7a and V1 in monkeys, were significantly positive (Figure 4 of (*1*)), indicating that task uncertainty is associated with lower perceptual confidence in both species.

### Recurrent neural networks trained on behavioural data generate mechanistic hypotheses

One challenge to understand the effect of task uncertainty is that, in natural environments and our task design, task uncertainty is inherently linked to negative feedback in the past. This relationship makes it difficult to distinguish responses to task uncertainty from innate biological reactions to the feedback.

To generate mechanistic hypotheses specifically about the cost of task uncertainty, we turned to a system without biological reactions to feedback. We trained recurrent neural networks (RNNs) to perform the same two-feature discrimination task and examined their internal dynamics. These models provide insights into potential neuronal mechanisms that can be validated through subsequent experiments.

We used two training strategies to create two distinct, comparable models. Both models received as inputs information about the stimuli on the current trial and the history of stimuli, choices, and rewards over the past nine trials. The models had a modular input-output structure, where the ‘task module’ received trial history inputs and was trained to output one of the two tasks, while the recurrently connected ‘perception module’ received stimulus inputs and was trained to output one of the four final choices (Fig. 2A; see Methods: *RNN models and analyses*). The modular structure was informed by evidence that task rule computations, which involve integrating evidence over several trials, are separable yet interact with sensory processing on a single trial (*1, 13*). Our goal was to generate hypotheses about how task certainty influences the computations underlying perceptual judgements on individual trials, so we focused on within-trial dynamics. We approximated the slowly varying trial history as a noisy but constant contextual input for the fast, transient stimulus inputs, whose time course was designed to match that of the monkey task. Note that all information about past trials must therefore be extracted by the models from the trial history inputs (see Fig. S9C-D for further analyses of trial history inputs).

**Fig. 2.**
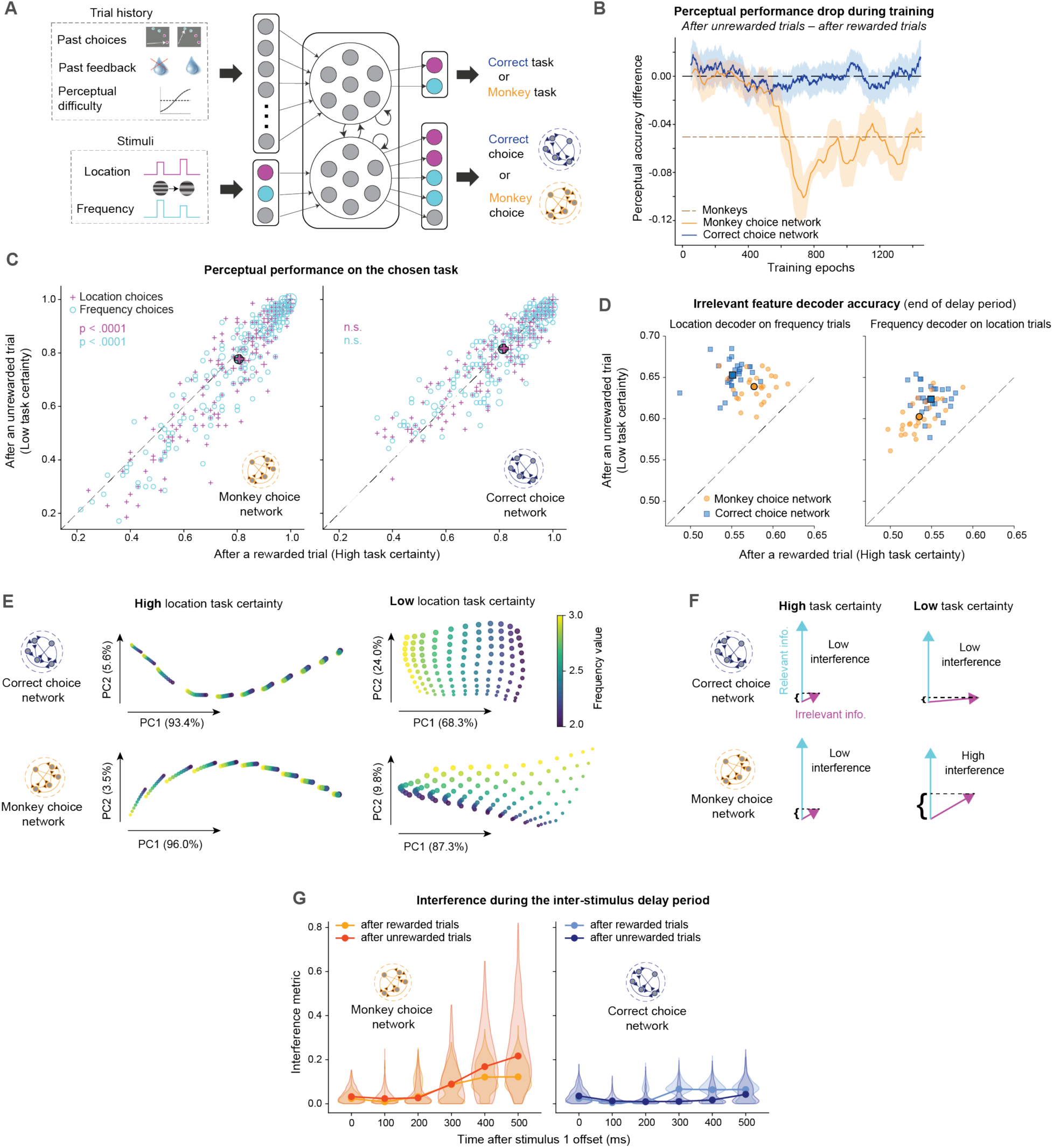
Generating mechanistic hypotheses using recurrent neural networks. (**A**) Model schematic for RNNs designed to either predict the correct choice (correct choice network) or the monkeys’ choice (monkey choice network) on each trial. Trial history inputs, which include the choices, reward feedback, and perceptual difficulty of the past nine trials, are fed into half of the recurrent units (top circled group, termed the “task module”). These units project to outputs that are trained to indicate the task on the current trial: location (magenta) or frequency (cyan). Current trial stimulus inputs, which include an input for location (magenta) and for frequency (cyan), are fed into the remaining half of the recurrent units (bottom circled group, termed the “perception module”) as two pulses separated by an inter-stimulus delay period. These units project to outputs that compute the choice on the current trial: location increase/decrease (magenta) or frequency increase/decrease (cyan). The task module and perception module are fully connected in the recurrent layer (short curved arrows). (**B**) The difference in perceptual accuracy (after unrewarded – after rewarded trials) throughout training for both models. The monkeys’ difference in perceptual accuracy was on average 5% (brown dashed line). The monkey choice network (orange) approaches this difference, while the correct choice network (blue) remains around zero Shaded regions show 95% confidence intervals from sliding window averaging (see Methods). (**C**) The monkey choice network (left) displays worse perceptual performance after an unrewarded, low task certainty trial than after a rewarded, high task certainty trial (p<0.0001 for both features, Wilcoxon signed-rank tests). In contrast, the correct choice network (right) did not show a significant difference in perceptual performance (p>0.05 for both features, Wilcoxon signed-rank tests). Each data point corresponds to a pair of accuracies (after a reward and after a nonreward) calculated from 70 trials, each with identical feature change amounts (stimulus condition). The marker size is proportional to the number of stimulus conditions that fall on that accuracy pair (most pairs correspond to one stimulus condition). (**D**) At the end of the delay period between stimulus presentations, information about the first stimulus irrelevant feature (location on trials where the model chose the frequency task (left) and frequency on trials where the model chose the location task (right)) can be decoded with higher accuracy from hidden layer activity of the perception module following unrewarded trials than following rewarded trials (p<0.0001 for both features and both models, Wilcoxon signed-rank tests). Each point corresponds to a pair of first stimulus feature values, which differ in the decoded feature value by a fixed amount and define the mean of noisy stimulus inputs to the model (see Methods). X- and y- axes show the performance of cross-validated support vector classifiers (after rewarded and after unrewarded trials respectively, 1000 trials each) in categorizing the two feature values. (**E**) Visualization of the RNN hidden layer representation of features of the first stimulus, 100ms before the onset of the second stimulus during the location task. Activation patterns were projected onto the first two principal components (>90% variance explained), which capture >90% of the variance. Each data point is one unique stimulus, with dot size corresponding to the location input, and color corresponding to the frequency input. Frequency (irrelevant feature) was better represented under task uncertainty (more separation between color groups after non-rewards than after rewards) for both networks. However, in the correct choice network, the relevant and irrelevant feature values vary along orthogonal directions, whereas in the monkey choice network the directions are less orthogonal, leading to feature interference. (**F**) A schematic of how stronger representations of the irrelevant feature when the task is uncertain combined with non-orthogonal representations of task variables leads to interference and thus a cost of flexibility. In an idealized correct choice network (top), the irrelevant feature is more strongly encoded (longer magenta vector) under low task certainty; however, it varies nearly orthogonally to the relevant feature, thus only leading to low interference. In the monkey choice network (bottom), the feature axes are more aligned, leading to high interference under low task certainty. The length of the vectors depicts the strength of the representations. (**G**) The monkey choice network, but not the correct choice network, has higher interference between the relevant and irrelevant stimulus features after unrewarded trials than after rewarded trials at the end of the inter-stimulus delay period. We quantified interference as the dot product of the two feature unit vectors scaled by the ratio of distance covariances (irrelevant/relevant; see Methods). Kernel density estimates of the distributions over trial history conditions are plotted at each timepoint along with the medians (2000 trial histories total at each timepoint). Distance covariances and angles are plotted separately in Figure S6.

The first model, the “correct choice network,” was trained to produce the correct choice given the stimulus feature values and its own trial history (Fig. 2A). Since, like our participants, it was trained to maximize rewards, the correct choice network offered a reference for comparison with humans and monkeys who are subject to the biological constraints in the brain. Like the ideal observer, the correct choice network shows no link between task uncertainty and perceptual performance at any point during training (Fig. 2B).

The second model, the “monkey choice network,” was trained to predict the monkeys’ actual choices given the stimulus feature values and the history of stimuli, choices, and rewards that the monkeys experienced before each trial (Fig. 2A). Indeed, the monkey choice network predicted a greater proportion of the monkeys’ choices, particularly errors, than the correct choice network (see Fig. S3 for these and other RNN performance metrics). Because of the way it was trained, unlike the correct choice network, the monkey choice network exhibited the reduction in perceptual performance after unrewarded trials characteristic of the monkey and human participants’ behaviour. In the model, this drop persisted from about halfway through training (Fig. 2B).

After training, the weights were frozen, and the monkeys’ trial history was used as the input to both models. Using the actual trial history, the correct choice model still did not exhibit worse perceptual performance under task uncertainty (Fig. 2C and S4B). This observation rules out the possibility that this cost arises solely from suboptimal choice history.

To complement and validate the comparison of perceptual performance on trials after a reward vs. no reward as a proxy for high vs. low task certainty (Fig. 2C), the modular structure of the networks allowed us to measure task certainty directly from the task module’s output activity, separately from the perceptual choice. Indeed, we found that this measure of task certainty varied as expected with feedback on preceding trials (Fig. S4A) and predicted perceptual performance (logistic regression; Fig. S4B).

### Task uncertainty leads to feature interference

The differences between our monkey choice network and correct choice network provide a platform for generating hypotheses about the mechanistic origins of the cost of task uncertainty. While the monkey choice network shows decreased perceptual performance under task uncertainty, the correct choice network flexibly switches between tasks without a cost to perceptual performance (Fig. 2B, 2C, S4B), much like the ideal observer (Fig. S1A). We reasoned that differences between the models could explain the cost of task uncertainty in humans and monkeys.

We compared the stimulus feature representations in the hidden layers of the two networks to understand why only the monkey choice network’s perceptual accuracy drops under low task certainty. We focus on the critical time point at the end of the inter-stimulus delay period, when any agent performing this task needs to compare the remembered features of the first stimulus with incoming sensory inputs about the second stimulus to inform the choice. One possibility is that the performance cost reflects a relationship between task uncertainty and encoding strength, such that the more relevant a feature is believed to be, the more strongly it is encoded in activity throughout the delay period (*2, 3, 5–7, 9, 10, 12*). Indeed, we observed this effect in the monkey choice model; however, we also observed it with equal (if not greater) effect size in the correct choice model. The believed irrelevant feature could be decoded with higher accuracy under low task certainty (after unrewarded trials) than under high task certainty (after rewarded trials) at the end of the delay period in both models (Fig. 2D). Looking at the relaxation dynamics following stimulus offset revealed that while both features were initially encoded with similar strengths, the believed relevant feature encoding was maintained throughout the delay period, while information about the irrelevant feature decayed at a rate dependent on task certainty in both models. (Fig. S5). Thus, encoding strength and random noise alone cannot explain why the monkey choice network exhibits the performance drop while the correct choice network does not.

However, comparing the geometry of the stimulus representations in the two models revealed important differences. Using distance covariance analysis (DCA) (41), we identified stimulus feature axes––directions in hidden unit space along which activity covaries the most with feature inputs––at different timepoints in the delay period (see Methods). We found that under low task certainty, when irrelevant information is enhanced, the two feature axes remained orthogonal throughout the delay period in the correct choice model but not in the monkey choice model (Fig. S6, angles bottom row, after unrewarded trials). These differences in geometric arrangement are also apparent when visualizing the stimulus representations in a (dimensionality reduced) space where the responses of each unit is one dimension (Fig. 2E, Fig. S8): in the correct choice model, the relevant and irrelevant feature values vary along orthogonal directions, whereas in the monkey choice model, they interfere. Thus, enhancement of irrelevant information did not affect the ability of the correct choice network to inform choices based on the relevant feature, as the representations of the two features remain independent. In contrast, in the monkey choice network, the non-orthogonal encoding combined with enhancement of irrelevant information when the task is uncertain leads to interference, where judgments of one feature (e.g., spatial frequency) are contaminated by information from the other (e.g., location).

In both models, under high task certainty, the DCA feature axes need not be orthogonal at the end of the delay period once the representation of the irrelevant feature has decayed (Fig. S6 angles bottom row, after rewarded trials). This highlights the interaction between encoding strength and orthogonality in producing interference; when irrelevant information is very weakly represented compared to relevant information, even non-orthogonal representations lead to little interference. In the extreme case, a feature that is not represented has no notion of direction in activity space.

We summarize the DCA results in Figure 2G with an interference metric defined as the dot product of the feature unit vectors scaled by ratio of distance covariances (irrelevant/relevant; see Methods). This measure of feature interference is consistent with the behavioural differences observed between the models. Example trajectories in principal component space are shown in Supplementary Figure S7, depicting how activity evolves towards different fixed points based on task certainty estimated from trial history inputs before stimulus onset (Fig. S7A), thus determining the relaxation dynamics back towards the fixed point after stimulus offset (Fig. S7B).

In summary, our analyses suggest that feature interference under uncertainty arises from two factors: (1) enhanced encoding of irrelevant information, observed in both networks; and (2) non-orthogonal neuronal representations of task-related features, observed only in the monkey choice network. Consequently, only the monkey choice network exhibited impaired perceptual performance under uncertainty, resembling human and monkey behaviour. We next tested this hypothesis using additional behavioural, neuronal, and causal experiments designed to characterize errors linked specifically to task uncertainty.

### Behavioural evidence for feature interference

We used behavioural methods to test the first prediction of our feature interference hypothesis: that irrelevant information is more strongly retained under low task certainty. Specifically, our monkey choice network predicted that during the delay, irrelevant feature encoding decays, but the decay is considerably slower under task uncertainty (Figure S5). Because we could not directly measure memory decay in monkeys, we conducted additional human psychophysics experiments. On 20% of the trials, we asked human participants to report one of the features (orientation or frequency) of the first stimulus after short (0.5 s) or long (2.0 s) delays (Fig. 3A top illustration). The participants’ ability to recall the believed irrelevant feature dropped substantially after long delays (p<0.0001, Fig. S1F), consistent with the prediction from the monkey choice network. This indicates that even though the RNN was trained on monkey behaviour, it can also predict aspects of human behaviour, thereby strengthening the case for homology between monkeys and humans and validating the RNN’s predictions. More importantly, consistent with the idea that irrelevant information is better retained in short-term memory when the task is more uncertain, participants had higher accuracy (smaller errors) when recalling the believed irrelevant feature on low task certainty trials (Fig. 3A magenta data points significantly below identity line, p=0.001). There was no significant change in relevant feature encoding (p=0.1, Fig. 3A, cyan points) from these feature recall trials.

**Fig. 3.**
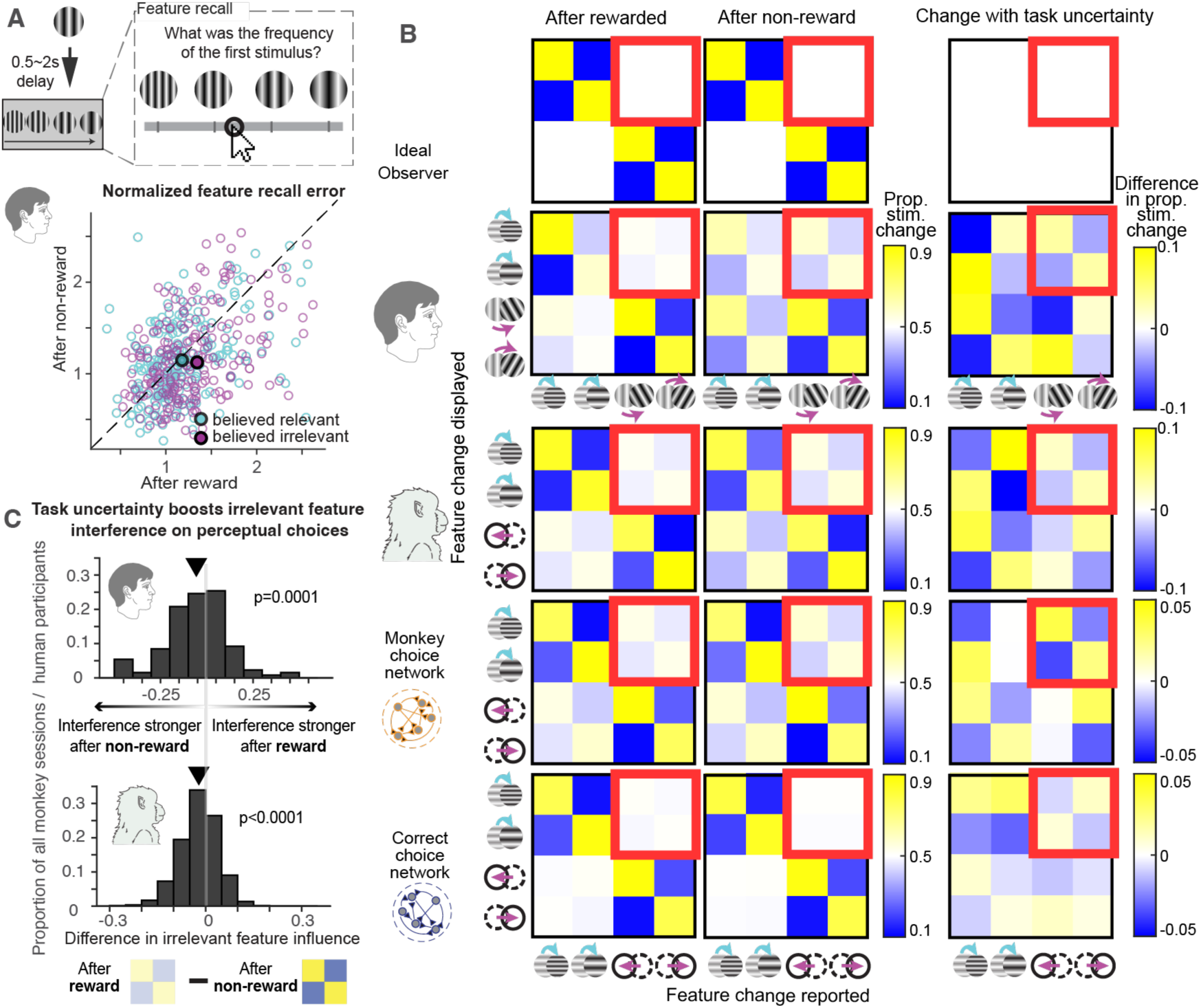
Behavioural evidence for the feature interference hypothesis. (**A**) Feature recall with human participants. In a subset of the trials, we asked human participants to reproduce one of the features (spatial frequency or orientation) of the first stimulus according to their memory following a delay. The illustration depicts the experimental design, including the slider used to reproduce the stimulus. In the scatter plot, each open symbol denotes each participant’s average error in the believed relevant (cyan) or irrelevant (magenta) feature under high task certainty (x-axis) and low task certainty (y-axis). The recall error for believed irrelevant feature was smaller for that feature after an unrewarded trial (low task certainty) than after a rewarded trial (high task certainty) (the magenta data points lie significantly below the identity line, p=0.0019). The participants’ ability to remember the believed relevant feature is not significantly different under low and high task certainty (p=0.2) (**B**) The relationship between choice and the irrelevant feature in an optimal observer (top), example human participant (second row), example monkey session (third row), the monkey choice network (fourth row), and the correct choice network (bottom). For each row, the three confusion matrices demonstrate the proportion of feature changes (rows in each matrix) that precedes each of the four choices (column in each matrix) under high task certainty (left matrices), low task certainty (middle matrices), and their difference (low-high, right matrices). Red squares highlight the portion relevant for feature interference. The choices of an ideal observer are unaffected by irrelevant features regardless of task certainty. Humans and monkeys exhibited patterns of dependence on the irrelevant feature that were stronger after unrewarded trials. The monkey choice network reproduced these error patterns, whereas the correct choice network, similar to the ideal observer, did not. (**C**) Perceptual choices are significantly more predictable from the irrelevant feature under lower task certainty. For each monkey experiment session and human participant, we predict the perceptual choices on one feature based on the changes in the other feature for trial-number matched conditions of low and high task certainty. Histograms (top: humans, bottom: monkeys) show that across monkey experimental sessions/human participants, the distributions of differences in prediction performances under high and low task certainty were significantly below zero (p=0.0002 for humans, p<0.0001 for monkeys), indicating that irrelevant feature is more predictive of perceptual choices under task uncertainty.

The enhanced encoding of the irrelevant feature might impair perceptual performance through either or both of two possible mechanisms:

1. The cognitive processing of the two features competes for limited attentional resources, so the irrelevant feature encoding under task uncertainty decreases attention to the relevant feature. This would predict more random perceptual errors under task uncertainty.
2. The neural representation of one feature systematically biases judgments of the other because their representations are non-orthogonal (feature interference), as in our monkey choice network. This would predict a systematic influence of the irrelevant feature on perceptual choices under uncertainty.

To distinguish these mechanisms, we analyzed the relationship between perceptual choices and the irrelevant feature in monkeys and humans. While the ideal observer does not show either the choice dependence on the irrelevant feature or a change in the dependence with task uncertainty (Fig. 3B, lack of a pattern in the highlighted parts of the confusion matrices in the top row), choices in both species were influenced more by the irrelevant feature following an unrewarded trial than following a rewarded trial. This can be seen as a stronger interference pattern under task uncertainty (the interference pattern in the highlighted portions of example confusion matrices on the second and third row of Fig. 3B grew stronger under task uncertainty.)

The enhanced interference patterns under task uncertainty in humans and monkeys were qualitatively captured by the monkey choice network (the highlighted portions of the fourth row of confusion matrices in Fig. 3B), but not by the correct choice network (the highlighted portions of the fifth row of confusion matrices). Across monkey sessions and human participants, the direction of irrelevant-feature influence was idiosyncratic (Fig. S2C–D). However, its magnitude was consistently greater under low task certainty within sessions. For each session, we quantified the proportion of choices predicted by the idiosyncratic association between choices and the irrelevant feature under low and high task certainty. Across species, choices were more strongly predicted by the irrelevant feature under low task certainty (Fig. 3C). This confirms that the increased perceptual errors under uncertainty reflect an amplification of specific interference between feature representations, and are not solely attributable to lack of attention to the relevant feature or the entire stimulus, or to increased randomness in the perceptual decisions.

### Neuronal evidence for feature interference

To test the neuronal predictions of our feature interference hypothesis, we recorded from area V1, as it robustly represents spatial frequency and location. We reasoned that V1 recordings would comprise a strong test of hypothesis, as V1 is not known for strong task-related modulations (*14, 15*). Thus, observing task uncertainty dependent interference in V1 suggests such interference likely exists in higher visual areas under dynamic task uncertainty conditions. For the following analyses, we focus on V1 responses to the first stimulus, since V1 activities are very low during the delay period without a stimulus in the receptive fields. Using a strategy similar to the one we used to identify feature interference in the activity of our modeled units, we first identified the linear combination of V1 responses that best encoded each visual feature (see Methods section *distance covariance analyses*). We tested the hypothesis that there is more feature interference under task uncertainty by analyzing both the signal and noise in the projections of V1 population responses on to the two feature axes.

To look for interference in the way the two features are encoded (signal), we attempted to decode stimulus frequency from the neural population axis that best encodes location, and vice versa. If the two features were encoded orthogonally, we would be unable to decode information about one feature from the axis representing the other. The model predicts that under task uncertainty, 1) the representations of the two features are non-orthogonal, and 2) the representation of the irrelevant feature will be enhanced (less variability and narrower distributions on the location axis in Fig. 4A lower panel than upper panel). Together, these factors will increase interference between the two features under task uncertainty.

**Fig. 4.**
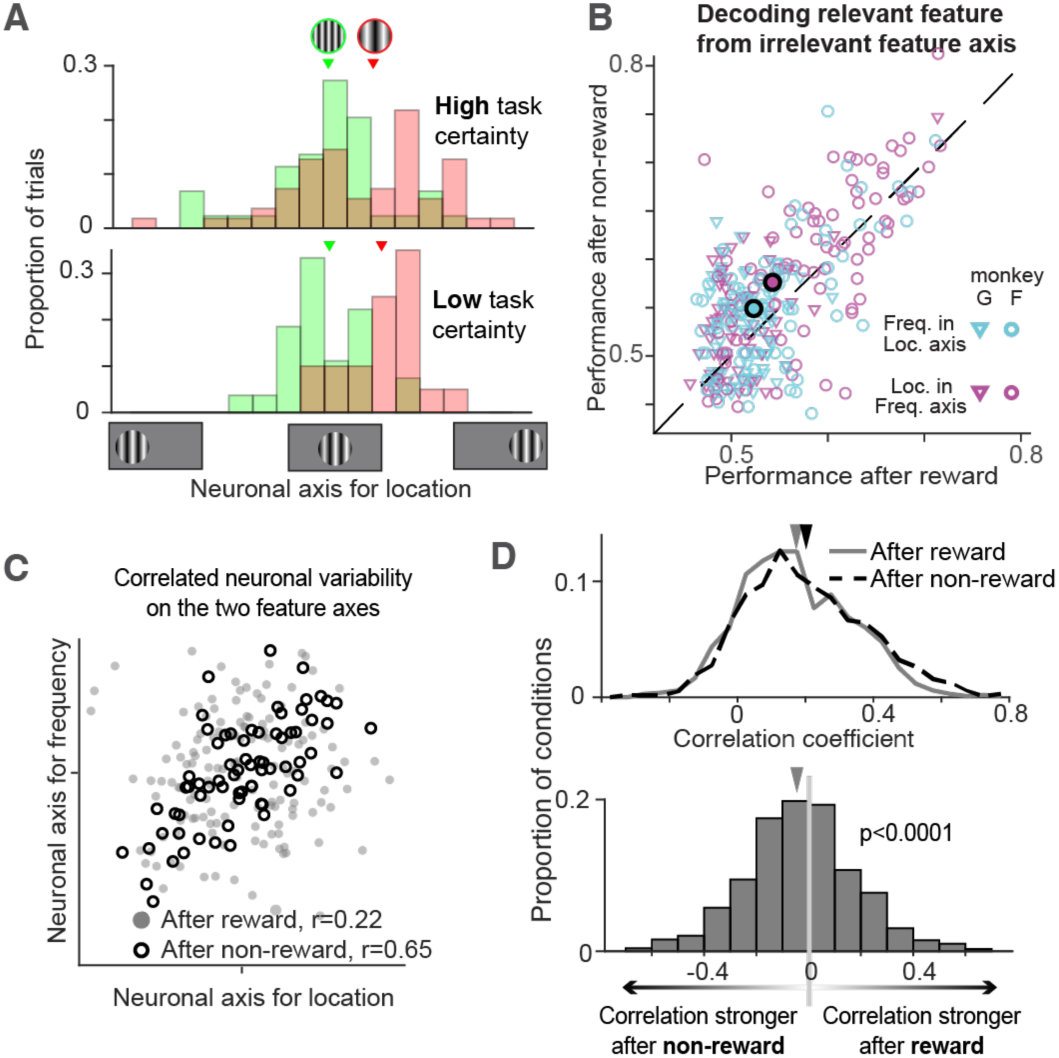
Neuronal and causal evidence for feature interference from monkey V1. (**A**) In an example condition with fixed stimulus location, stimuli with different frequency values lead to more distinctive distributions of V1 response projections (red and green histograms) onto the location encoding axis under low task certainty (bottom) than high task certainty (top). For both task certainty condition, we carried out leave one out cross-validation to quantify decoding performance. (**B**) Decoding performances across all stimulus conditions in all sessions. For both features, the points are distributed significantly above the identity line (p=0.006 for location classifiers in frequency dimension, p=0.036 for frequency classifiers in location dimension), indicating increased feature interference under low task certainty. (**C**) Trial to trial variability of V1 population responses to an example stimulus in an example session projected onto the spatial frequency and location axes. These projections are more correlated under low task certainty (black circles) than under high task certainty (grey dots), indicating stronger interference under task uncertainty. As there is no a priori prediction about the sign of the correlation, positive correlation is simply defined as the overall correlation between projections on the two feature axes in the session. Correlation coefficients are calculated for matched number of trials in high and low task certainty. (**D**) Correlation coefficients for all stimulus conditions in all sessions. The upper panel shows the overall distributions for correlation coefficients under high task certainty (gray solid line, median correlation 0.16), and low task certainty (black dashed line, median correlation 0.18). The two distributions are significantly different (Wilcoxon rank sum test, p=0.001). Pairwise differences in the coefficients for the same stimulus (as in C) also showed that the correlations are significantly stronger under low task certainty than under high task certainty (lower panel, p<0.0001), again demonstrating stronger feature interference under task uncertainty.

We used linear discriminant analysis to find the encoding axis for both features in V1 population response space. Consistent with the feature interference hypothesis, across all stimulus conditions and sessions, we found that cross-decoding performance was significantly enhanced under task uncertainty (Fig. 4B both cyan and magenta data points lie significantly above the identity line).

To look for evidence for feature interference in trial-to-trial variability (noise), we analyzed the relationships between the projections of trial-to-trial variability of V1 population responses to repeated presentations of the same stimulus onto the two feature axes. The model predicts that the two feature axes are not orthogonal, which would lead to correlations between the fluctuations of projections of population responses onto the two axes (Fig. 2F), and the correlation should be stronger under low than high task certainty (example stimulus condition in Fig. 4C). Consistent with this idea, we found that across conditions and sessions, the correlation between projections onto spatial location and frequency axes were more correlated under low task certainty (Fig. 4D), indicating stronger interference. Together, these task uncertainty dependent changes in feature encoding and trial-to-trial fluctuations in V1 are not expected if the perceptual drop with task uncertainty were trivially attributed to general confusion or inattention. They revealed the surprising possibility that task-related modulations of the interference between visual feature encoding happens on the sensory level as early as V1.

### Causal evidence for feature interference

Finally, we used a causal manipulation to test the prediction of our feature interference hypothesis that feature representations are less independent under task uncertainty. To generate specific, testable predictions, we injected current into each unit in the hidden layer of the monkey choice network. Like V1 neurons, RNN units also encode both location and frequency, so manipulating one unit affects behavior in both tasks. (Fig. 5A). We quantified the effect of this perturbation as shifts in choices on trials when the stimuli did not change (red arrows, Fig. 5A). Under high task certainty (after rewarded trials), perturbation-induced changes in choice behaviour were uncorrelated between tasks (Fig. 5B left). In contrast, under low task certainty (after unrewarded trials), these effects were significantly correlated (Fig. 5B) and significantly different from the high task certainty condition. This correlation of model perturbation effects between left-shift and frequency increase is also consistent with the direction of behavioral interference (Fig. 3B confusion matrices for monkey choice network). This generates the testable prediction that under low task certainty, when stimulation induces left-location shift reports, it will induce more frequency increase reports, and vice versa.

**Figure 5:**
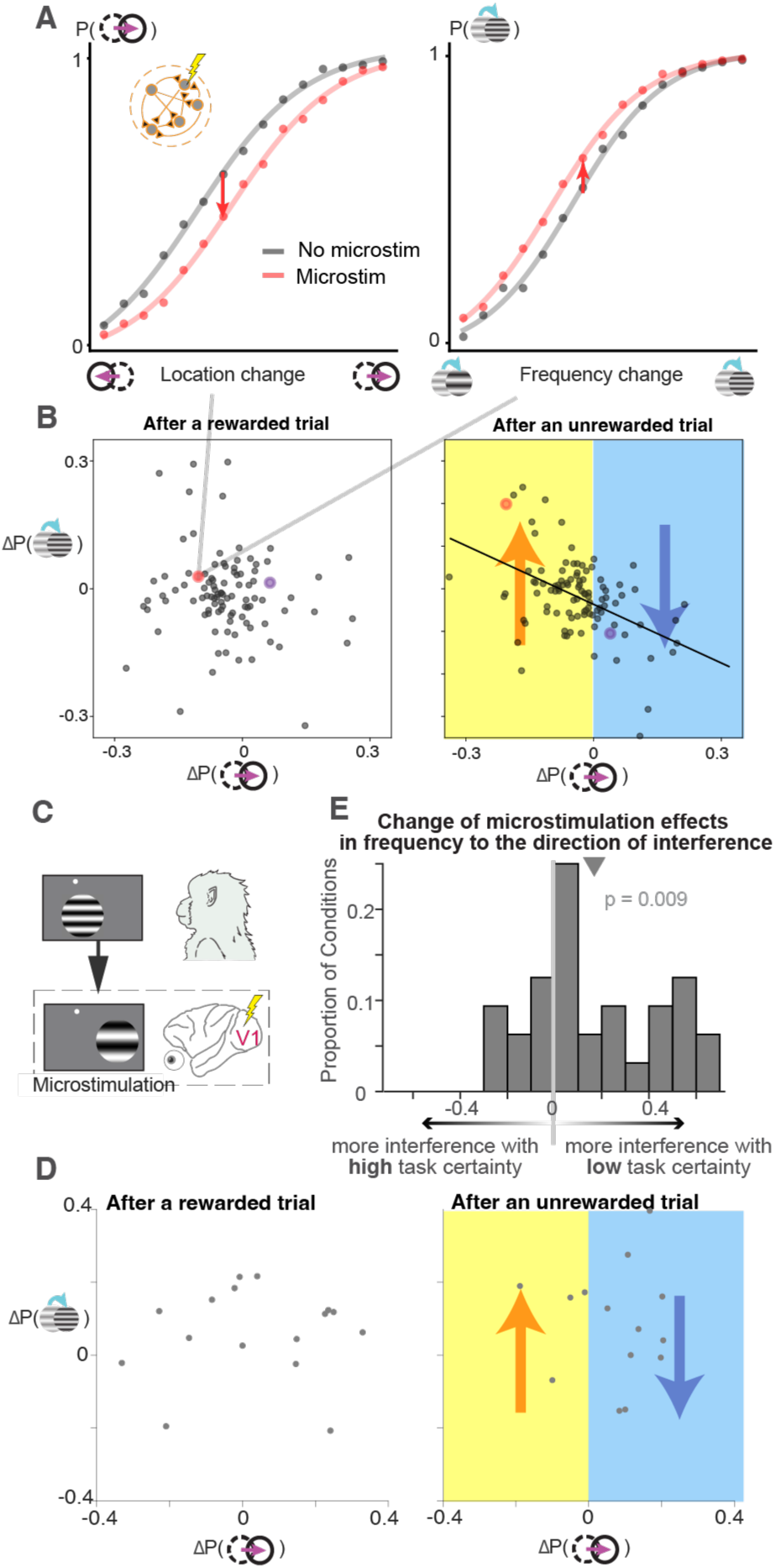
Causal evidence for feature interference. (A) Causal manipulations in the monkey choice network. We injected current into individual hidden units (“microstimulation”) and measured effects on psychometric curves for both tasks. Black curves are sigmoidal fits of the network’s original location (left) and frequency (right) psychometric curve. Red curves show the shifted psychometric curves when we perturbed the activity of one example model unit. We summarize the impact of perturbing a unit as the change in choices when there is zero stimulus change (red arrows); (B) Network perturbation effects across perceptual units under high and low task certainty. Perturbation effects were uncorrelated between tasks following a rewarded trial (left panel; Pearson r = –0.15, p = 0.2); but significantly correlated following an unrewarded trial (right; r = –0.33, p = 0.0007), which is also significantly different from that following the rewarded trial (p < 0.0001, bootstrapping). This predicts that task uncertainty changes microstimulation effects in the direction of the feature interference (red and blue arrows); (C) Electrical microstimulation of V1 at the onset of the second visual stimulus in one monkey. (D) Similar to (B), microstimulation effects from the same V1 site and stimulus conditions were compared under high (left) and low (right) task certainty. (E) Increased feature entanglement under task uncertainty. The histogram shows that the microstimulation effects in frequency task moved to the direction predicted by feature interference under task uncertainty (as shown by the red and blue arrows in (D), p=0.009).

To test these predictions, we picked one V1 electrode each session and electrically microstimulated at the onset of the second stimulus on a randomly selected half of the trials (Fig. 5C). Although it is difficult to characterize the exact groups of neurons activated by microstimulation (*16*), our goal was to measure the behavioral effects of microstimulation in both tasks (Fig. 5D, similarly calculated as Fig. 5A-B for RNN perturbations), and compare them under high and low task certainty (Fig. 5D left vs. right). Assuming that stimulation impacted a similar ensemble of V1 neurons throughout the session, systematic differences in behavioral outcomes between low- and high-certainty conditions should reflect differences in how those features are represented. Consistent with this prediction, we observed that microstimulation effects were more entangled under task uncertainty. By comparing microstimulation effects between high and low task certainty (Fig. 5D), we confirmed that the latter demonstrated more feature interference: under task uncertainty, on trials where the monkey did the frequency task, microstimulation biased choices in the direction predicted by the model (Fig. 5E), indicating that the representations of location and spatial frequency were more entangled.

## Discussion

Humans and animals must flexibly switch between tasks in ambiguous contexts to survive in an uncertain world. This flexibility has long been known to come at a cost. We used a multidisciplinary approach to identify neural reasons for the cost associated with switching between tasks. By grounding our investigation in neural population-level representations of tasks and corresponding visual features, we uncovered a neuronal feature interference mechanism that unifies previous findings and has implications for our understanding of many cognitive processes.

### Feature interference as a neuronal mechanism for task uncertainty costs

We aimed to uncover the neuronal mechanisms underlying the behavioural cost of flexibility observed in cognitive psychology studies (*2–12*). Various hypotheses have been proposed to explain slower, poorer choices during task switching. These include limits on processing capacity and undue attention to irrelevant information, inhibition of relevant signals, task-set reconfiguration, and proactive interference. Here, we demonstrate that the task uncertainty cost could be the combined result of two factors: partially overlapping (entangled) feature representations and enhanced retention of information that is irrelevant to the chosen task under task uncertainty.

These findings align with previous theories proposing that after a task switch, proactive interference from the pre-switch task hinders adaptation to the new task (*10, 17*). Our findings extend these insights by demonstrating that task uncertainty, even without actual task switches, can induce feature interference and that this interference can be linked directly to entangled representations in populations of neurons throughout the brain.

Our findings suggest that while the extent of feature coupling systematically increases with task uncertainty, the direction of coupling is idiosyncratic across subjects and even experimental sessions in our monkey subjects. The origins of these baseline associations between independent features remain unclear. They may reflect peripheral constraints on frequency discriminability, slow fluctuations in neuronal variability over hours or days, or individual differences shaped by life experience, experimental context, or correlations in natural environmental statistics. While our study focused on the neuronal mechanisms and behavioral consequences of increased interference under uncertainty, understanding the sources of these idiosyncratic associations is an important direction for future work that includes normative accounts and developmental perspectives.

### Integrative approach reveals neuronal mechanism for complex behaviour

The key insights in our study were enabled by the comprehensive approach combining artificial neural networks, monkey electrophysiology, and human psychophysics, to understand the neuronal mechanism for advanced cognition and complex behaviour. Classic uses of artificial neural networks in neuroscience research involve training them to produce correct choices on laboratory tasks, in the hope that the trained networks reveal the computations that occur in the brains of participants performing the same tasks (*18, 19*). Here, we trained our models to mimic actual behavioural responses including error patterns (*20, 21*) and generated testable neural hypotheses, which we further tested using large-scale electrophysiology and causal manipulations in monkeys. Human psychophysics further complemented the animal work, extending the relevance of our findings across species and ensuring applicability to broader cognitive phenomena. Combined, our approach holds the potential to extract meaningful insights from complex data, a critical challenge in an era when datasets are becoming increasingly large and complex (*22*).

### Feature interference in a network of brain regions

We observed neural signatures of feature interference even in V1, a primary sensory area in which neuronal activity is minimally affected by cognitive processes (*14, 15*). This suggests that the phenomenon of interfering neural representations is not confined to specific brain regions but rather extends throughout the brain.

Our results suggest several promising avenues for future work. For example, it is plausible that feature interference poses an even stronger limit in higher-order visual areas and prefrontal cortex, which are more intimately linked with advanced cognitive processing. Indeed, studies have suggested that orthogonal neural representations in prefrontal cortex enable context-dependent decision-making, with mixed results about the gating of irrelevant features (*23, 24*) .

Our monkey choice network makes one prediction that is not testable in V1 because it does not have prolonged delay period activity: that feature interference grows stronger over time during the delay period (Fig. 2E and Fig. S6). An important question for future work is whether other brain areas with persistent representations of visual stimuli, such as prefrontal cortex, exhibit growing feature interference over time as stimulus information is held in working memory.

Additionally, attention is known to reduce perceptual and neural interference from irrelevant stimulus features and objects (*25–27*). While our results indicate that the task uncertainty cost cannot be attributed solely to temporary inattention, feature interference might be implemented by attention-like gain modulations on neurons that jointly represent relevant and irrelevant features in working memory. Future work should investigate the origin and temporal dynamics of this interference. The effect could arise from the neuronal circuitry alternatively sampling between feature dimensions under uncertainty (*28*). Furthermore, our previous work demonstrated surprising neuronal interactions between task-related information decoded from macaque area 7a and perceptual signals decoded from V1 (*1*). This raises important questions: are feature entanglements in V1 directly caused by parietal (or other top-down) feedback? What might be the functional role of these inputs to visual cortex, if they seem to limit feature representation under task uncertainty? Answering these questions will be crucial for understanding the constraints for flexible decision-making in health and disease.

### Dimensionality as a fundamental limit on cognition

Our findings, together with a broad body of earlier work, suggest potential neural mechanisms underlying feature interference. One clue comes from studies showing that cognitive processes such as attention reshape population activity by altering its dimensionality, partly through changes in the balance of excitation and inhibition (*29–32*) . A reduction in dimensionality could cause representations of distinct features to overlap, producing the interference observed here. Although participants were required to process only a single stimulus with two relevant features on each trial, inferring the hidden task required tracking recent stimuli, choices, and rewards demands that substantially expand the dimensionality of the required state representation. Reports from human participants and the behavior of monkeys indicate that the task strained working memory, consistent with the idea that interference may emerge when representational dimensionality approaches capacity.

These observations raise the broader possibility that the dimensionality of neural population activity constitutes a fundamental limit on cognition. Because dimensionality determines how many variables—features, objects, task states, or timescales—can be represented independently, it constrains the behavioral computations a population can support. Although dimensionality could theoretically scale with population size, empirical estimates are typically far lower (*33, 34*), aligning with well-known capacity limits across working memory (*35*), attention (*36*), decision-making (*37*), and learning (*38*).

Computational models suggest that low-dimensional representations enhance generalization but sacrifice discriminability (*39, 40*). However, it remains unclear under what biological constraints low-dimensional representations naturally arise in task-driven circuits. Future work using recurrent neural networks with more realistic top-down modulation and synaptic plasticity may help clarify how networks balance representations across multiple timescales. Biophysically realistic constraints on models that encourage dimensionality reduction in complex natural environments may also explain the origin feature entanglement in simplified laboratory tasks.

Evidence across species is broadly consistent with dimensionality limits on cognition. Forming associations between simple features (e.g., color and texture) that jointly predict a complex feature can reduce dimensionality (*41*); undoing these associations may require expanding dimensionality, potentially explaining difficulties in reversal learning (*42*), rule learning in bipolar disorder (*43*), and language learning (*44*). Relatedly, the typical working memory capacity around five items (*35, 45*) may reflect a limit on independent neural dimensions, as strategies such as chunking effectively compress representations into fewer dimensions (*18, 45*). These converging lines of evidence suggest that diverse cognitive limitations may share a common neuronal origin.

### Translational implications of feature interference

Deficits in cognitive flexibility are an especially stark consequence of the many brain disorders, including dementia, substance abuse, and developmental disorders, that are associated with diminished cognitive capacity (*46, 47*). If feature interference indeed underlies many cognitive limitations, our results offer hope for improving and repairing cognitive capacity. For example, we recently demonstrated that methylphenidate (trade name Ritalin, a stimulant used to treat disorders of attention) increases the dimensionality of neural population representations in the visual cortex and that this increase is correlated with improvement in a visual task (*48*). This is a proof of principle supporting the idea that repurposing existing drugs to address other cognitive abilities that might share a neural mechanism with their primary target could be an effective way to improve cognition in health and disease. The suggestion in our V1 results that feature interference affects brain-wide mechanisms means that systemic drugs and/or therapies that affect widespread neuromodulatory systems might be especially effective.

Together, our results hold promising implications for the future of basic science and translational efforts to mechanistically understand, improve, and repair cognition. Our feature interference hypothesis provides a concrete set of testable predictions with implications for theoretical and experimental work spanning many species and systems.

## Methods

### Behavioural task for non-human primates

We designed a two-interval, two-feature discrimination task with stochastic task switching that allows us to measure and manipulate task certainty and perceptual decision-making (*3*). Briefly, two rhesus monkeys (both male, 12 and 9 kg) fixated a central spot while two Gabor stimuli (∼2° visual angle in diameter) were displayed in series (200 ms each), separated by a variable delay (300-500 ms). The second Gabor stimulus differed from the first in its spatial location (shifted left or right) and spatial frequency (higher or lower), with independently randomized change amounts in the two features. After a 150 ms second delay, the fixation dot disappeared, and the animals looked at one of four peripheral targets to indicate both the inferred relevant feature and the direction of change in that feature (spatial frequency increase or decrease and spatial location left-shift or right-shift). Perceptual performance refers to the proportion of correct perceptual discriminations of whichever feature the monkey chose to discriminate. Small titrations of the change amounts in the two features were made before each experiment session based on the animals’ performance in the previous session, so as to roughly keep the perceptual performance of the animals in both tasks around 75%. The actual across-session average perceptual performance is 79% for monkeys. Each point in Fig. 1D depicts the perceptual performance for the stimulus condition with the same relevant feature change amount for an entire recording session. The median number of trials per condition was 233. The relevant feature changed stochastically on 2.5% of trials, and the monkeys were rewarded only if they correctly discriminated the relevant feature. The visual display contained no information about the behavioural relevance of features, so the animals needed to infer the relevant feature based on their stimulus, choice, and reward history. The median number of trials in each daily session was 1086. All animal procedures were approved by the Institutional Animal Care and Use Committees of the University of Pittsburgh and Carnegie Mellon University, where the animal experiments were conducted.

### Behavioural task for the human participants

We adapted our task for online human psychophysics (built with Gorilla experiment building tools (*49*) and customized scripts). We recruited a total of 206 adult human participants (97 female, 109 male) through Prolific (prolific.co), who performed slightly different combinations of behavioural experiments. All experiments were performed in accordance with the IRB protocol at Carnegie Mellon University. Informed consent was obtained from all human research participants.

In the main experiments, we used a version of the two-interval, two-feature discrimination task with stochastic uncued task switching that we adapted to better suit human participants and online experiments (Fig. 1B). The timing was slightly different (500 ms initial fixation period, 300 ms stimulus presentations, and 500 ms delay period). The two features to be discriminated were orientation and spatial frequency. As in the monkey experiments, the changes in the two features were independently varied. Similar to the monkeys, the change amounts were set at each participant’s perceptual threshold, which was found through a 3-up/1-down psychophysical staircase procedure before the main task session. Participants indicated their choice by clicking on one of four buttons representing each feature and its change direction (indicated by a combination of text and graphical icon). As in the monkey experiments, the human participants received feedback following each choice (a green tick indicated a correct report of the direction of change in the task-relevant feature, and a red cross indicated an incorrect perceptual judgment, task choice, or both). As in the monkey experiments, the relevant feature switched stochastically, but the frequency was different (with a probability of 0.1 on each trial for humans, 0.025 for monkeys). The overall length of a typical session varied between 100 to 150 trials and around 15-20 minutes.

### Task-belief and feature perception confidence report

Some task sessions required participants to report their confidence in their perceptual judgment on a random 20% of trials. After reporting their decision on the main task but before receiving feedback, participants were presented with three confidence sliders. Participants were instructed to indicate their task-belief on the first slider, ranging from a strong belief that spatial frequency was relevant to a strong belief that orientation was relevant. An intermediate value indicated low task certainty. The second and third sliders were used to report the perceived direction of change for each feature, where the absolute value reflected confidence in the perceptual judgment.

### Variable delay and feature recall question

In some sessions, we explored memory for believed relevant and believed irrelevant stimulus features by interrupting a randomly selected 20% of trials after either 500 ms (short delay) or 2000 ms (long delay) to ask participants to recall a feature of the first Gabor. On each trial, each feature value was chosen from a fixed set of seven evenly spaced values (orientation from 0° to 54° in steps of 9°; at a distance of 53cm, frequency range from 2 cycle/° to 3.35 cycle/° in ratios of 1.09). Participants indicated their feature memory using a slider, facilitated by four reference Gabor patches that spanned the possible range. We quantified recall error as the difference between the actual stimulus feature and the recalled feature. For each feature, we normalized the participants’ recall error by the difference between neighbouring feature values (Fig. 3A, Fig. S2B), so an error of 1.0 would represent 9° in orientation, and an 9% increase in frequency.

### Recurrent neural network models and analyses

#### Training and architecture

We trained recurrent neural network (RNN) models using custom code based on PsychRNN (*50*) and TensorFlow (*51*). The models were governed by the following dynamical equations:

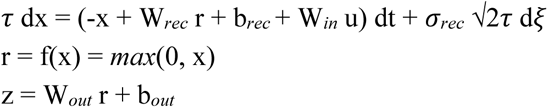

where *x* is the recurrent state, which when passed through the rectified linear unit (ReLU) activation function gives the firing rate *r*, *u* is the input vector, and *z* is the output vector. *W_in_*, *W_rec_*, and *W_out_* are the input, recurrent, and output weight matrices, and *b_rec_* and *b_out_* are constant biases added to the recurrent and output units. *dt* is the discrete time step, 𝜏 is the neuronal time constant, and *𝜎_rec_* scales recurrent noise, which is a Gaussian noise process (d𝜉). Both RNNs had *𝜎_rec_* =0.1, 𝜏=100, and dt=10. The RNNs were trained using supervised learning with a mean squared error loss function (target outputs described below). We used L2 regularization on weights (all coefficients 0.03) and firing rates (coefficient 0.06). Optimization was performed using the Adam algorithm (learning rate 0.003).

### Inputs and outputs

Our monkey choice network (Fig. 2A) was trained to predict the choices of two macaques based on the same history of stimuli, choices, and rewards experienced by our monkey participants. The RNN had two modules, which we term the ‘task module’ and the ‘perception module.’ This modular structure was implemented to separate the processing of slow trial-history inputs from the fast, transient stimulus inputs and to aid interpretability. We verified that this architectural choice was not the primary driver of our results; a fully connected (non-modular) network produced qualitatively similar results, including the behavioral drop (Fig. S9A) and the increase in feature interference (Fig. S9B) in the monkey choice network but not the correct choice network. The two modules were fully connected in the recurrent layer and trained simultaneously, but received different inputs and mapped onto different outputs. The task module received a 54D trial history input vector encoding the monkeys’ choices (one-hot encoded 4D vectors), stimulus feature change amounts (for the chosen task, signaling perceptual difficulty), and rewards (+1 for reward, -1 for no reward) for nine preceding trials. These inputs were constant throughout the trial, based on the assumption that task rule dynamics are slow compared to single-trial stimulus response dynamics. The task module was trained to output the task that the monkey chose on the current trial as a one-hot encoded 2D target output vector. We experimented with the number of preceding trials (range 5-12), and within this range we did not see substantial effects on the model’s ability to predict the monkeys’ choices, although predictably, decreasing this number to two trials of history did decrease performance. Nine was chosen as a conservative number within this range, as we did not want to limit performance based on trial history. Analysis of network performance as a function of history length showed that the model’s predictive power plateaued after 3-5 trials (Fig. S9C). Further, the network trained with a 9-trial window inherently learned to assign the most weight to the most recent trials, largely ignoring inputs from trials further in the past (Fig. S9D).

Meanwhile, following a 50 timestep initial delay period, the perception module received two transient stimulus inputs––2D feature vectors representing the spatial frequency (SF) and spatial location (SL) of the stimuli––lasting 20 timesteps each and separated by a variable delay period (30 to 50 timesteps, uniform distribution). The first stimulus vector had values uniformly distributed between 2 and 3. The second stimulus vector was generated such that its difference from the first was proportional to the actual frequency and location changes presented to the monkey on that trial. Throughout the stimulus presentations, the perception module also received a “fixation input” equal to 1, indicating a target output of zeros (no response). 15 timesteps after the second stimulus presentation, the fixation input fell to 0, indicating the choice period. The perception module was trained to output the monkey’s choice, one of the four choices (SF increase, SF decrease, SL decrease, SL increase) as a one-hot encoded 4D target output vector. There was one additional output trained to mimic the fixation input. This time course was designed to model the monkey task, with 1 RNN timestep representing 10 ms. Units in the perception module displayed mixed selectivity for the two features.

Gaussian noise with zero mean and variance 2* 𝜎*_in_ ^2^* was added to the inputs. 𝜎*_in_* was set to 0.5 for the trial history inputs to the task module and 0.2 for the fixation input to the perception module. For the stimulus inputs to the perception module, we chose 𝜎*_in_* =0.8 such that the perceptual accuracy of the monkey choice model matched that of the monkeys. We then used the same parameters for the correct choice model described below.

### Correct choice model

We compared the monkey choice network to a correct choice network (Fig. 2A), which was trained to produce the correct choice based on the perceptual inputs of the current trial and its own trial history, with feedback provided based on the model’s choices rather than the monkeys’ choices. The correct choice network shares the same architecture and input structure as the monkey choice network to allow direct comparison. Stimulus change amounts were drawn from the same data used to train the monkey choice network. It is important to note that the correct choice network is, therefore, different from the network reported in our previous study (*1*). Here, we focus on the neural dynamics within each trial under varying task uncertainty conditions, leveraging new data, models, and analyses to investigate the neural mechanisms behind the perceptual performance drops under task uncertainty that we observed previously. Thus, the network models presented here are specifically designed to explore within-trial dynamics, such as feature interference. This differs from the model in our previous publication, which focused on dynamics across trials.

### Analyses

The correct choice network perceptual accuracies during training in figure 2B were calculated directly from its training batches (200 trials per epoch) and smoothed using a sliding window average over 100 epochs. The monkey choice network perceptual accuracies were calculated post-hoc by simulating 2000 trials per condition (after reward, after non-reward) using weights saved every 15 epochs during training, and smoothed using a sliding window average over 6 datapoints (90 epochs). For the remaining analyses, we chose to analyze the monkey choice network at the point during training with the highest monkey choice accuracy (Fig. S1), which was at training epoch 1245 out of 1500. We also used the monkeys’ trial history inputs for both models after training so that we could directly compare trials (all model figures besides Fig. 2B). This also allows us to rule out the possibility that the effect of task uncertainty on perceptual performance comes only from the monkeys’ suboptimal choice history, in which case the correct choice model should exhibit this effect as well when given the monkeys’ trial history inputs. For the single-trial PCA trajectories, input and recurrent noise was set to zero (Fig. S7). For all remaining analyses we used the same noise levels as during training, except for the DCA feature axis analysis noise to the stimulus inputs was reduced slightly from 𝜎*_in_* =0.8 to 0.5.

### Distance covariance analysis for assessing orthogonality of feature axes

To quantify the orthogonality of feature axes in the models, we used distance covariance analysis (DCA), a dimensionality reduction method that identifies linear projections that capture potentially nonlinear interactions between variables by maximizing the correlational statistic distance covariance (*52*). The method aims to find the covariance between distances in stimulus feature space and distances in the neuronal population response space. Specifically, for the stimulus feature matrix *Y*, DCA aims to find the best neuronal response subspace that maximize the squared distance covariance:

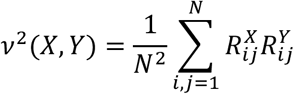

Where *N* is the number of samples, and 𝑅^X^ and 𝑅^X^ are the distance matrices in neuronal space and stimulus space (both recentered so that the row, column, and matrix means are all zeroes). By examining model activity with continuously varying stimulus inputs to the model, DCA reveals the neuronal activity axis that covary the most with each feature values, whether they covary linearly and nonlinearly. After reducing the dimensionality of the activity of all units in the hidden layer to the first 10 principal components (capturing >95% of the variance), we calculated the distance covariance statistic and corresponding axis separately for each unique set of trial history inputs, feature (location and spatial frequency), and timepoint during the inter-stimulus delay period, from 500 simulated trials with continuously varying feature values. We define the interference metric (*IM*; Fig. 2E) as the dot product of the two feature unit vectors scaled by the ratio of the corresponding distance covariances:

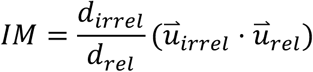

For equivalent analysis in the neuronal data, we analyzed V1 responses to the initial stimulus of each trial, which typically had only two possible location values and two possible frequency values per session. Under these conditions, DCA is effectively equivalent to linear discriminant analysis. Therefore, for simplicity, we used linear discriminant analysis on the spike counts of V1 in response to the initial stimulus in Figure 4. We applied linear discriminant analysis on the responses of the recorded V1 neurons to find the encoding axis for either feature in neural response space. To quantify feature interference in the neuronal population representations, we quantified our ability to decode one stimulus feature on the best axis for decoding the other feature.

### Ideal Observer Model

We employed a normative model similar to our previous work (*1*) to simulate an ideal observer with perceptual abilities matching those of the participant. Using the participant’s psychometric curve and the stimulus change magnitude on each trial, we estimated the probability of an incorrect perceptual choice. For every non-rewarded trial, the likelihood odds that the rewarded task is different from vs. the same as the previous trial grows by:

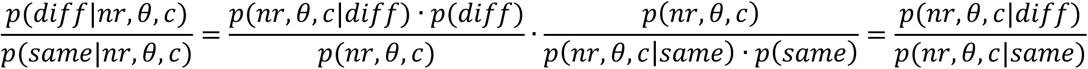

where 𝜃 and c denote the stimulus change and perceptual choice, and nr indicates a non-reward trial. We assumed an overall equal number of trials with the relevant feature as either one of the two (i.e., any two randomly picked trials from a session are equally likely to have the same task or have different tasks, or p(same)=p(diff)). Because participants were never rewarded when their belief was incorrect, p(nr|diff)=1. When the belief matches the task rule, the perceptual error probability is derived from the corresponding psychometric function

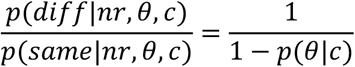

For n consecutive non-reward trials, the likelihood ratio favoring a task switch grows as shown in

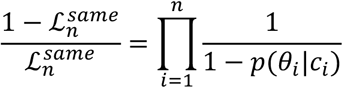

while the prior probability that the task remains unchanged after n non-reward trials is given by

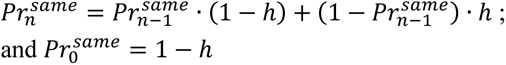

with hazard rate h=0.025 for monkeys, and h=0.1 for humans. Combining perceptual evidence with prior volatility, the model predicts a task switch when the posterior probability of the task being the same with the previously chosen is smaller than that of the other task.

Switch to the other task when: 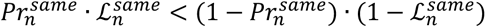

Finally, once the task is determined, a random choice between feature change directions is generated according to the probability informed by the participant’s psychometric curve.

### Electrophysiology for the monkey experiments

Different aspects of a subset of the electrophysiological data have been reported previously (*3*). The visual stimuli were displayed on a linearized CRT monitor (1,024 × 768 pixels, 120-Hz refresh rate) placed 57 cm from the animal. We monitored eye position using an infrared eye tracker (Eyelink 1000, SR Research) and used custom software (written in MATLAB using the Psychophysics Toolbox, (25) to present stimuli and monitor behaviour. We recorded eye position and pupil diameter (1,000 samples per s), neuronal responses (30,000 samples per s), and the signal from a photodiode to align neuronal responses to stimulus presentation times (30,000 samples per s) using hardware from Ripple. We recorded neuronal activity from chronically implanted Utah arrays (Blackrock Microsystems; 48 electrodes in V1) during daily experimental sessions for several months in each animal (90 sessions from monkey F and 68 sessions from monkey G). We set the threshold for each channel at three times the standard deviation and used threshold crossings as the multiunit activity on that unit. We positioned the stimuli to maximize the overlap between potential stimulus locations and the joint receptive fields of V1 units. The receptive fields were measured using separate mapping sessions, during which the monkeys fixated their gaze on a dot, while Gabor stimuli were subsequently flashed at random locations for 100ms each across a portion of the screen.

We included experimental sessions for analysis if the total number of completed trials was at least 480 (which was ten times the number of channels on one multielectrode array). During complete trials the monkeys successfully maintained fixation until they indicated their choice. In Fig. 4, we analyzed the activity of V1 units during the initial stimulus display period, shifted by 34 ms neuronal response latency (*53*). V1 responses were excluded from analyses when a microsaccade was detected during this time window. V1 units are included for analyses if the stimulus response is 25% larger than baseline activity, measured 100 ms before stimulus onset, and 2) larger than 5 sp/s. These procedures resulted in 89 sessions from Monkey F and 65 sessions from Monkey G; an average 46 V1 units, and a median 1086 completed trials per session. Other aspects of these data have been reported in a previous study (*1*).

We did V1 microstimulation experiments during 6 sessions in one monkey (monkey F, Fig. 4). We were unable to conduct these experiments in monkey G because the experiments were interrupted by the COVID shutdown of the lab, after which there were no usable channels on the array. During microstimulation sessions, we electrically stimulated one channel on the V1 array with 20, 30 μA, 200 Hz biphasic pulses between 50 and 150 ms after the onset of the second Gabor stimulus on a randomly selected 50% of trials (*16*).

Although previous studies demonstrate that microstimulation activates a sparse, spatially distributed population of neurons whose axons run very near the electrode tip (*16*), we cannot directly infer the anatomical clustering of feature-selective neurons activated by microstimulation in our study. However, this lack of knowledge of the anatomical distribution of the stimulated neurons does not affect these results. We used this causal manipulation to identify differences in the effect of microstimulation under low and high task certainty. Assuming that the group of neurons whose responses are modulated by microstimulation does not depend on different task certainty, systematic differences in behavioral outcomes between low and high task certainty indicate that task uncertainty alters the functional coupling between features.

### Statistical tests

All p-values reported in this study are from Wilcoxon signed rank tests unless otherwise specified. We used linear mixed effect model (*54*) to account for random factors in samples coming from the same sessions (e.g. for the performance differences in Fig.1D-E). Within each session, we apply hierarchical bootstrap (*55*) of the trials to ensure balanced trial counts at the level of the participants’ chosen task, low and high task certainty, and each stimulus condition. Bonferroni correction was applied to all the statistical tests (e.g. across features and monkeys) related to the same hypothesis.

## Acknowledgments

We are grateful to John Maunsell, Douglas Ruff, Ramanujan Srinath, and Pouya Bashivan for comments on this manuscript.

This work was supported by the Simons Foundation (Simons Collaboration on the Global Brain award 542961SPI to MRC) and the National Eye Institute of the National Institutes of Health (awards R01EY022930, R01EY034723, and RF1NS121913 to MRC).

## Author contributions

C.X. and M.R.C. designed research; S.K.M. and C.X. trained recurrent neural networks and analyzed the data; C.X. and L.E.K. performed monkey training and neuronal recording; C.X. analyzed the neuronal data and performed the microstimulation experiments; R.C. and C.X. performed the online human psychophysics experiments and data analyses; M.R.C. supervised the findings of this work; C.X., S.K.M and M.R.C. wrote the paper.

## Competing interests

Authors declare that they have no competing interests.

## Data and materials availability

Code for recurrent neural network training and analysis is available at https://github.com/solkm/task-interference.

## Supplementary Figures

**Supplementary Figure S1:**
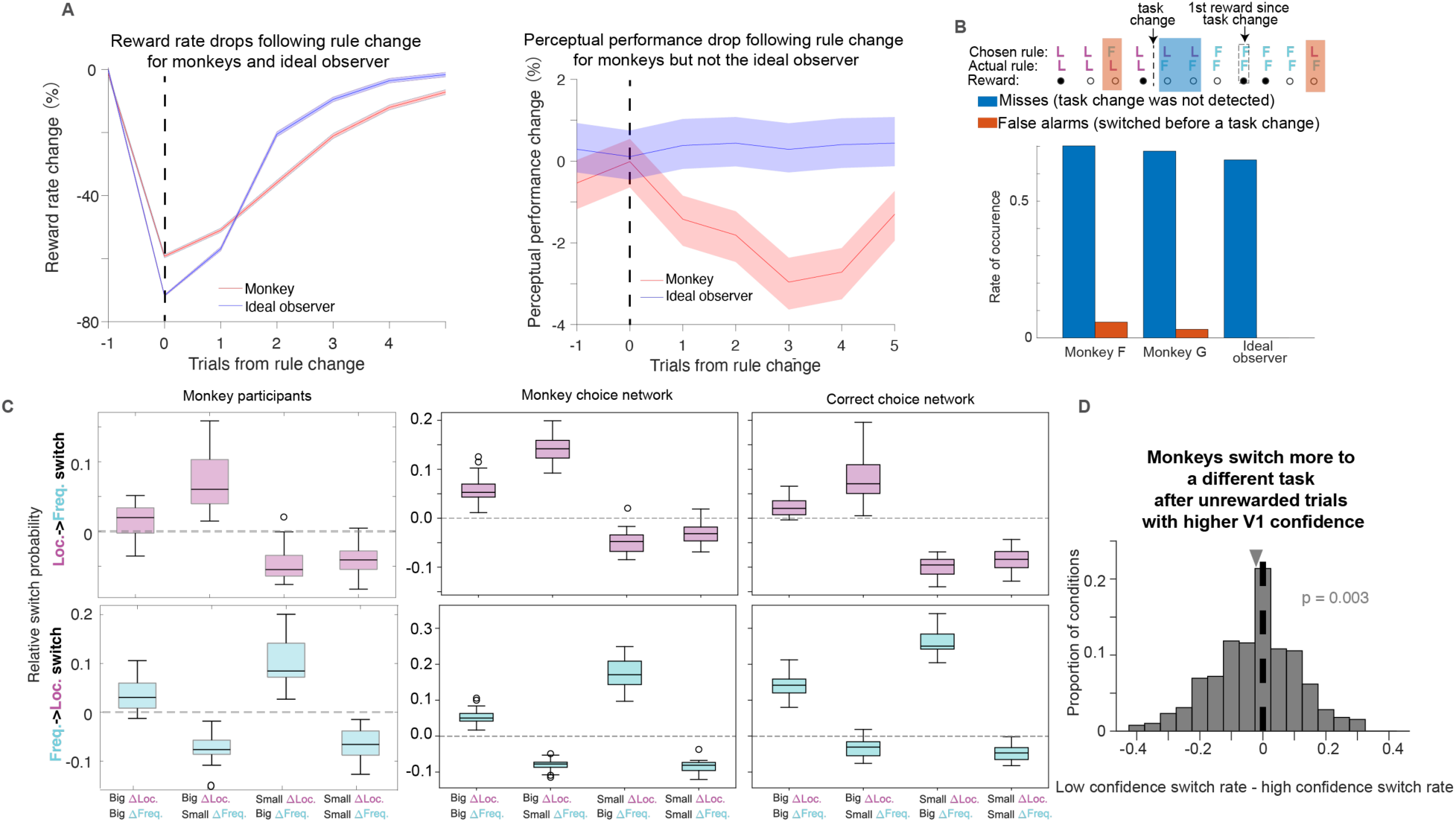
Task switching behaviour. (**A**) Reward rate (left) and perceptual performance (right) aligned to task-rule changes, for monkeys (red) and the ideal observer (blue), respectively. Shadings indicate s.e.m. (**B**) Types of task errors. Since a reward would validate the participant’s choice of task, any task errors that occurred between a task change and the first reward after that task change are considered “misses”; and any task error that occurred between the first reward after a task change and the next task change are considered “false alarms”. The rate of "miss" errors was substantially higher than that of "false alarms" for both monkeys, which is qualitatively similar to the ideal observer. Note that the “false alarm” rate for the ideal observer is 0.04%, not zero. (**C**) Switch rate after an unrewarded trials with different stimulus change amounts for monkeys (left), monkey choice network (middle), and correct choice network (right), relative to the average switch rate across stimulus conditions. To quantify the variability in these data, we divided the behavioral data into 20 non-overlapping subsamples. In both models and in the monkeys, the switch was higher following unrewarded trials featuring large, obvious changes in the believed relevant feature (first and second box in the upper plots, first and third box in the lower plots). (**D**) Switch rate difference after strong perceptual confidence non-rewards, and after weak perceptual confidence non-rewards, as measured by the distance of V1 population activity to the feature change hyperplane (Fig. 1B bottom). Even among identical unrewarded trials, stronger perceptual confidence decoded from V1 predicted more frequent task switches (histogram significantly to the left of 0).

**Supplementary Figure S2:**
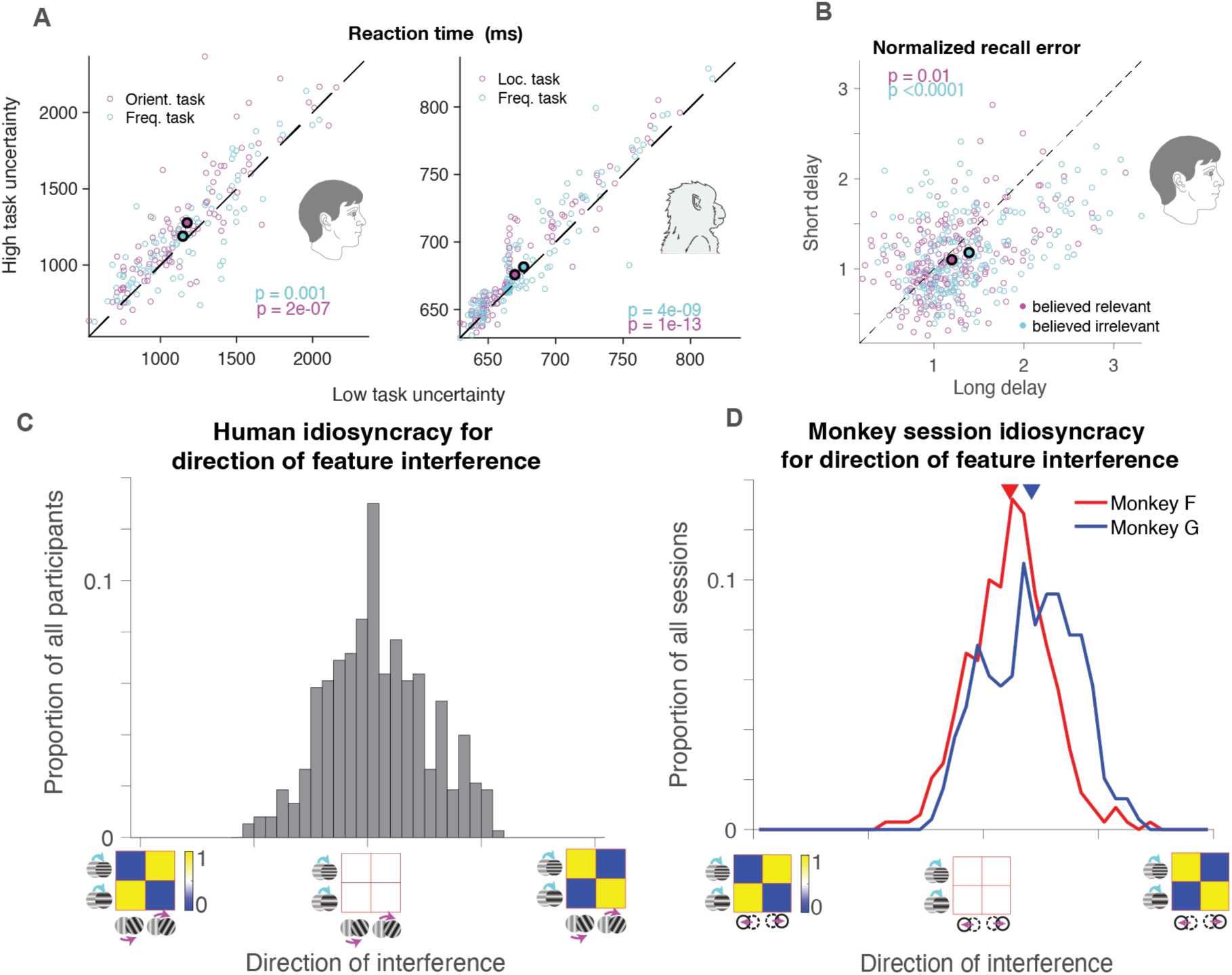
Additional participant behaviour. (**A**) Reaction times. Under higher task uncertainty, both humans and monkeys exhibited slower response times. Fig.1F is generated by taking the difference between the x and y axis of data points in this figure. (**B**) Recall error of features considered relevant vs. irrelevant. Each open symbol denotes each participant’s average error in the believed relevant (magenta) or irrelevant (cyan) feature after long delays (2 seconds, x-axis) and short delays (0.5 second, y-axis). Participants made smaller recall errors for both features after short delays than after long delays. (**C**) Heterogenous, idiosyncratic directions of feature interference across participants. For each human participant, we quantified coupling direction as the difference between the probabilities of “frequency-increase with counterclockwise-tilt” and “frequency-increase with clockwise-tilt” associations. The distribution across participants revealed diverse coupling directions. (**D**) Similarly, monkey sessions showed across session variability in the direction of feature interference, though both monkeys exhibited an overall “frequency-increase with location-left-shift” association.

**Supplementary Figure S3:**
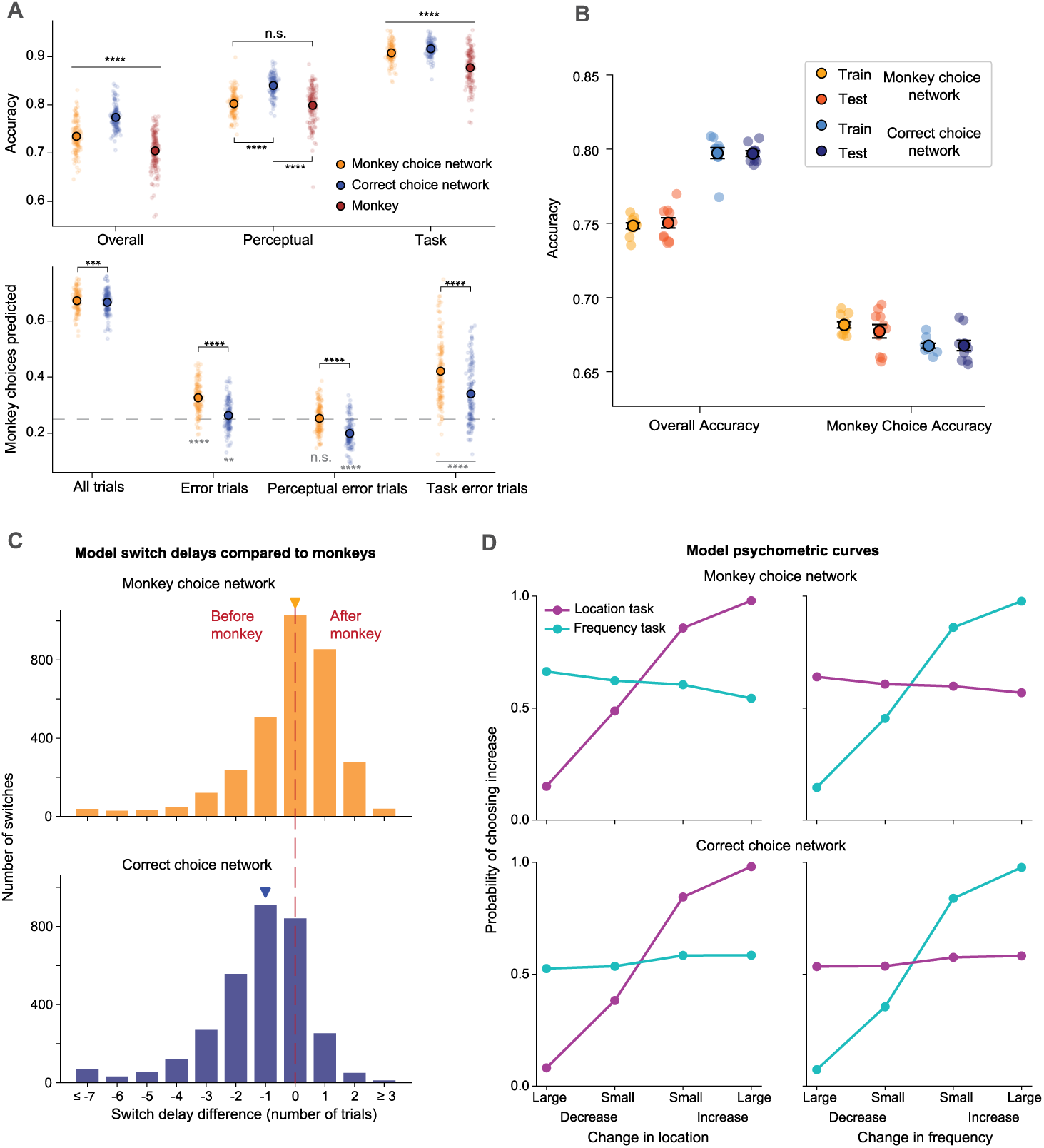
RNN performance and additional behavior. **(A)** Top: Accuracies of the monkey choice network (orange) and correct choice network (blue) after training, with comparison to the monkeys (brown). Overall: percentage of correct choices (chance=25%); Perceptual: percentage of correct perceptual judgments, regardless of the chosen task (chance=50%); Task: percentage of correct task choices, regardless of the perceptual judgment (chance=50%). Each data point corresponds to a session of monkey data, and outlined points indicate means. Bottom: Proportion of model choices that match the monkeys’ choice on the same trial (chance=25%, grey dashed line), shown for all trials and the subset of trials where the monkey made any error, a perceptual error, or a task error. Monkey task: percentage of task choices that match the monkeys’ chosen task, regardless of the perceptual judgment (chance=50%). Black annotations indicate comparisons between the two models, and grey annotations indicate comparisons to chance level. All statistical comparisons are by Wilcoxon signed-rank tests.. **(B)** Generalization performance of the monkey choice and correct choice networks on training data and held-out test data. We trained 10 models each with different random seeds, which determine the trials selected for training and testing. Each datapoint is a model. Solid, black-outlined points are the average over the 10 models, with black error bars indicating S.E.M. **(C)** The network models’ delay in switching tasks after a task change, relative to the monkeys’ delay. This is computed for trials where both the model and the monkey are correctly performing the previous task at the time of the switch. Triangle markers show the median switch delay difference. On average, the monkey choice network switches with the same delay as the monkeys. In contrast, the correct choice network switches one trial earlier on average. Note when the models switch after the monkey (delay difference > 0), the monkeys’ switch is included in the trial history inputs. **(D)** Psychometric curves of the network models for the location and frequency tasks. Both models learn to largely ignore the believed irrelevant feature (though the monkey choice network shows some signs of interference), with choice accuracy varying with perceptual difficulty (believed relevant feature change amount).

**Supplementary Figure S4:**
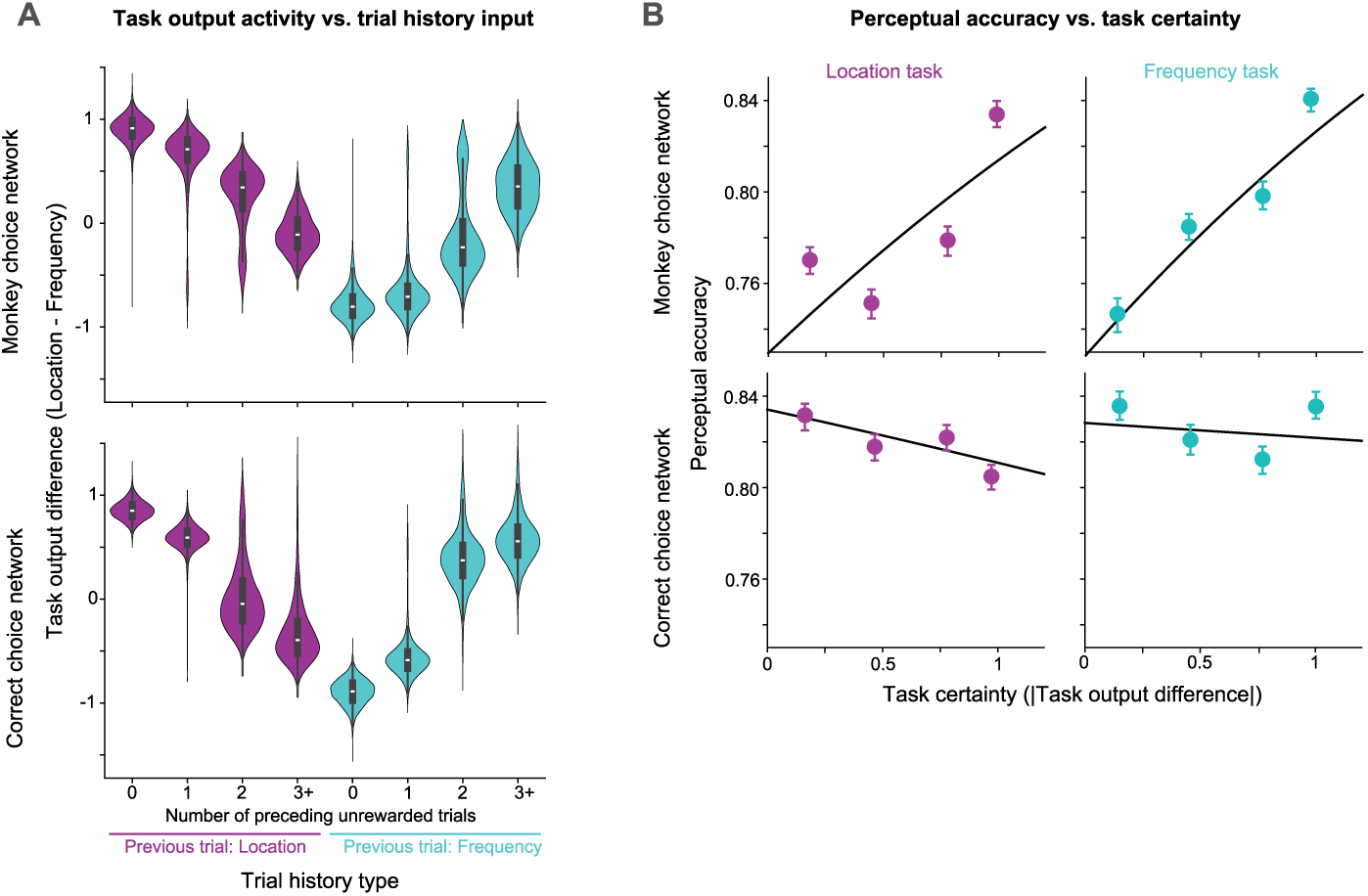
Task output activity provides a continuous measure of network task certainty, and perceptual accuracy increases with task certainty in the monkey choice but not the correct choice network. **(A)** Task output activity not only indicates the believed relevant task, but also a measure of certainty in both networks. “Task output difference” (y-axis) refers to the difference between the two output unit activities of the task module (location – frequency) at the last timepoint of the trial. During training, the target output was [1, 0] or [0, 1] when the target task was location or frequency, respectively (see Methods). Thus, a task output difference of +1 indicates a certain location choice, -1 indicates a certain frequency choice, and 0 indicates complete uncertainty. Indeed, in both networks, after a rewarded trial (0 preceding unrewarded trials on the x-axis) the task output difference is +1 or -1 depending on the previous trial task. As the number of non-rewards increases, the task output difference decreases or increases towards 0. Note that in the correct choice network, which tends to switch tasks with less delay (see Fig. S3C), the zero crossing happens after fewer non-rewards than in the monkey choice network. **(B)** Perceptual accuracy increases with task certainty in the monkey choice network (top row) but not the correct choice network (bottom row) for both tasks. Task certainty (x-axis) is defined as the magnitude of the task output difference described above. Scatter points indicate perceptual accuracy in four bins of task certainty (bins [0, 0.3), [0.3, 0.6), [0.6, 0.9), ≥0.9; x-value indicates mean). Error bars indicate a 95% confidence interval from 100 bootstrap samples. Black curves indicate logistic regression fits to the original datapoints (before binning). The coefficient for task certainty was positive for both tasks in the monkey choice network (location: b = 0.4842, 95% CI [0.440, 0.528], p<0.0001; frequency: b = 0.5736, 95% CI [0.529, 0.618], p<0.0001). In the correct choice network, task certainty had a small negative effect on perceptual accuracy for the location task (b = -0.1595, 95% CI [-0.215, -0.104], p<0.0001) and no significant effect for the frequency task (b = -0.0450, 95% CI [-0.092, 0.002], p=0.06).

**Supplementary Figure S5:**
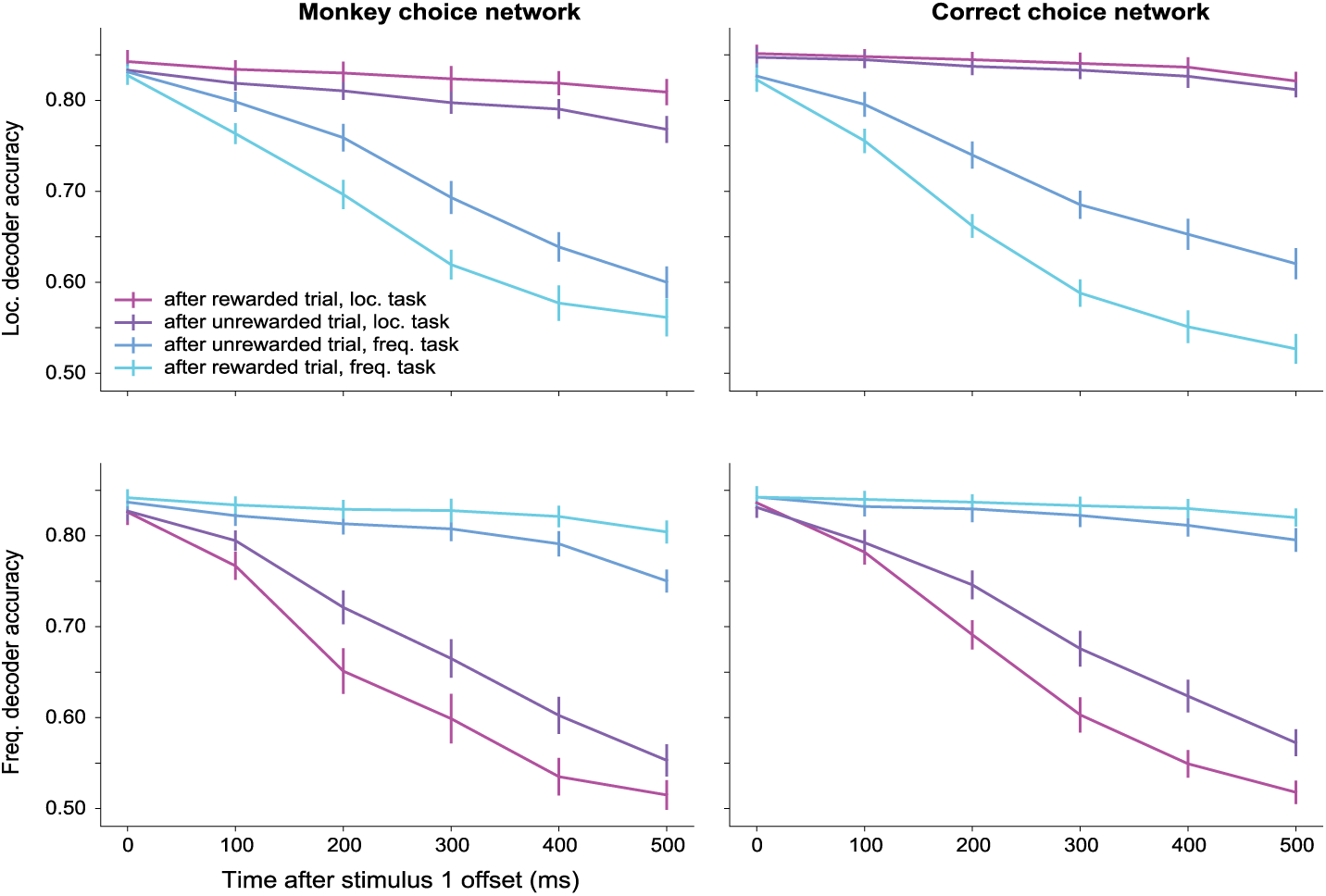
Feature (location, top row, and frequency, bottom row) decoder accuracies during a 500 ms interstimulus delay period for the monkey choice network (left column) and correct choice network (right column) in four trial conditions. The conditions are as follows: 1) after a rewarded trial, location task chosen (high location task certainty); 2) after an unrewarded trial, location task chosen (low location task certainty); 3) after an unrewarded trial, frequency task chosen (low frequency task certainty); 4) after a rewarded trial, frequency task chosen (high frequency task certainty). In both models, the more a feature is believed to be irrelevant, the more quickly information about that feature decays following stimulus offset. Each point represents the mean performance of cross-validated linear decoders (same as Fig. 2E, which shows performance at 400 ms), with error bars showing +/- 1 standard deviation. A separate decoder was trained for each stimulus condition (identical first stimulus) and timepoint.

**Supplementary Figure S6:**
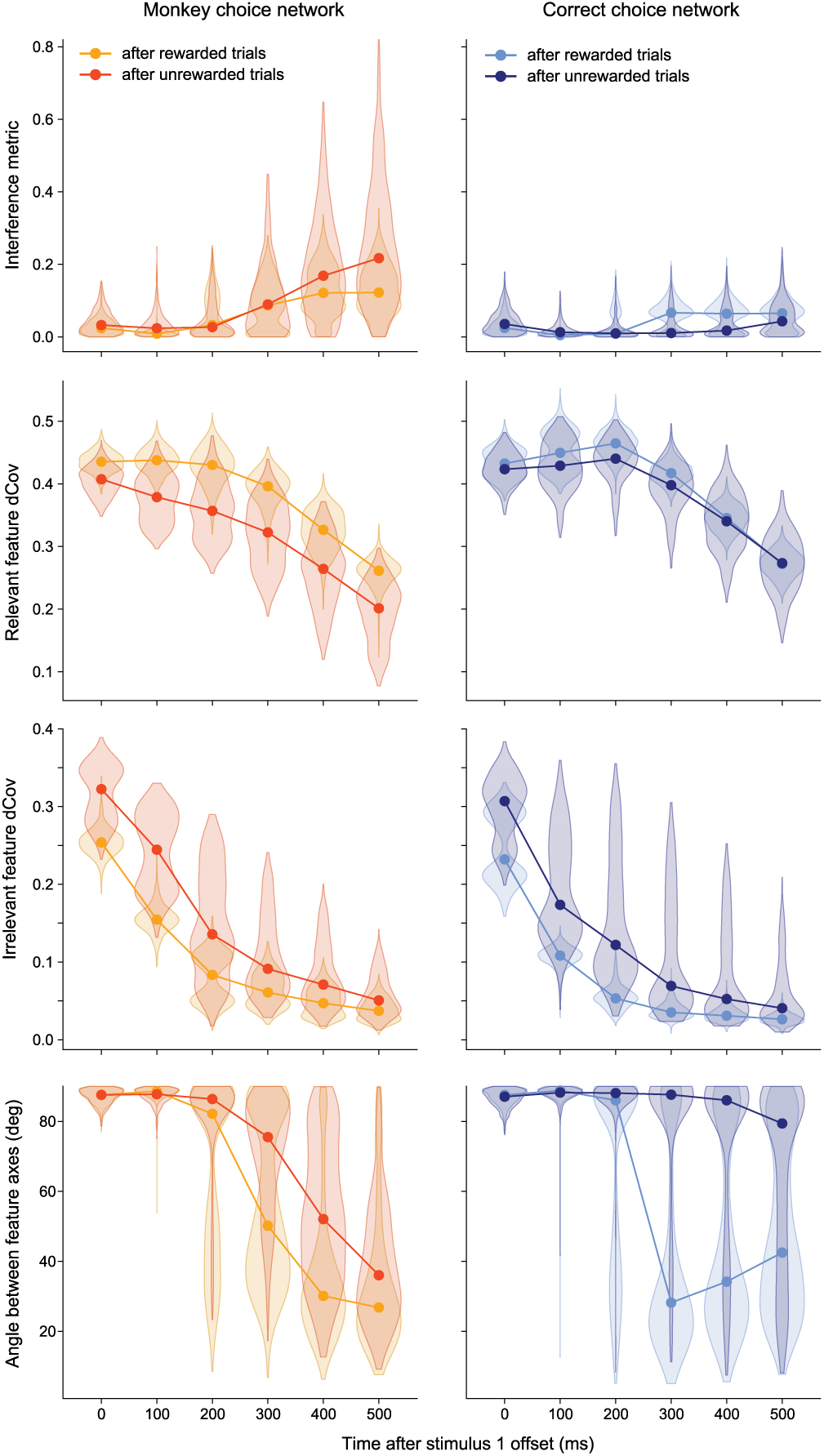
Distance covariance analysis to quantify feature interference in the monkey choice (left) and correct choice (right) networks after rewarded and unrewarded trials during the inter-stimulus delay period. First row: same as figure 2E. Second row: the distance covariance of the relevant feature is smaller after unrewarded trials in the monkey choice network throughout the delay period, and in the correct choice network at the start of the delay period. Third row: the distance covariance of the irrelevant feature is larger after unrewarded trials in both networks, throughout the delay period. Bottom row: angle between the feature axes, from 0 to 90 degrees (unsigned). While both networks exhibit a loss of orthogonality as the irrelevant feature representation decays after rewarded trials, orthogonality is maintained in the correct choice network, but not the monkey choice network, after unrewarded trials when the irrelevant feature is more strongly represented.

**Supplementary Figure S7:**
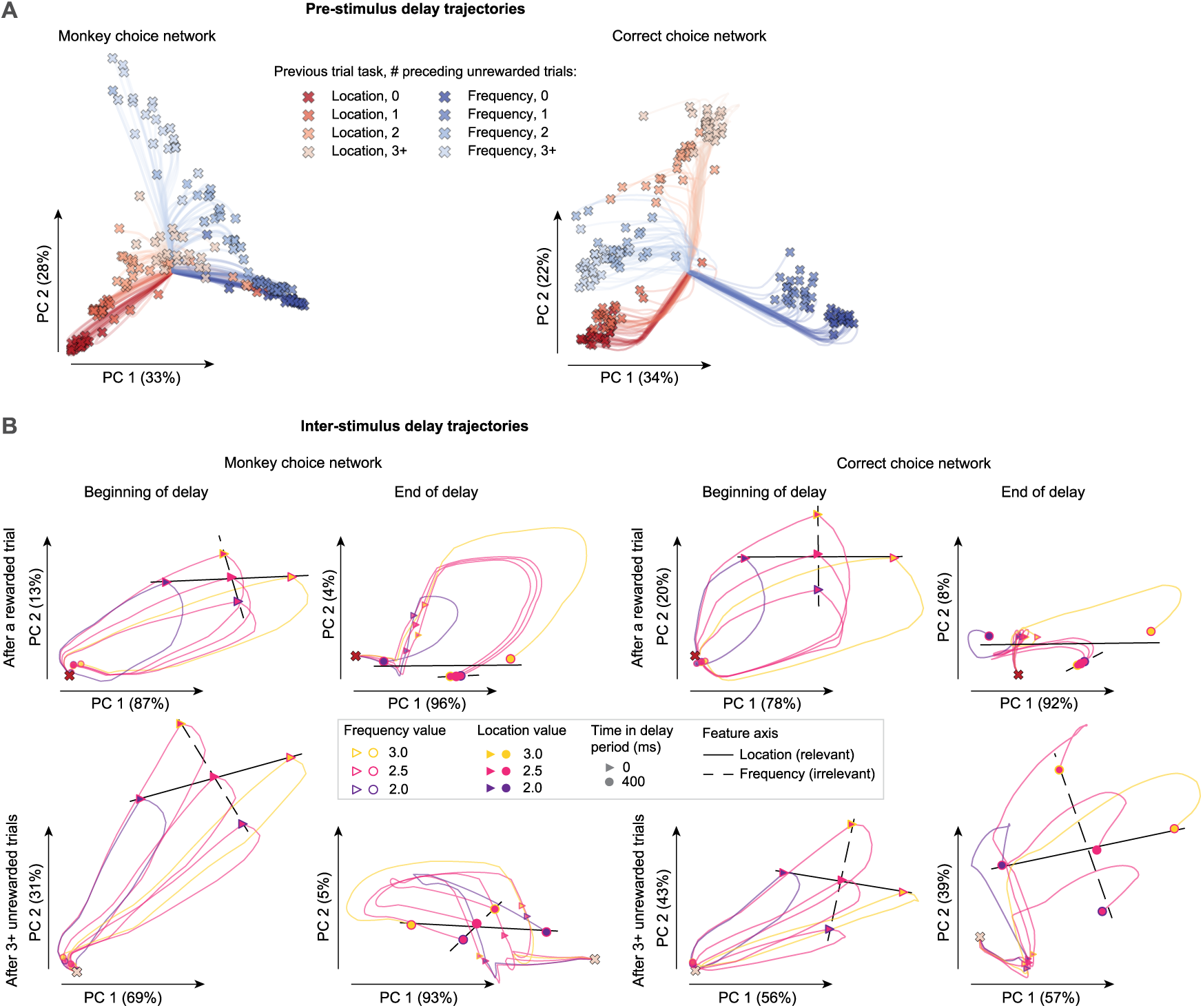
Single-trial trajectories in principal component space of the monkey choice and correct choice networks’ hidden layer activity. **(A)** Before stimulus onset, activity evolves towards fixed points determined by task certainty estimated from the trial history inputs, both in the monkey choice network (left) and the correct choice network (right). Trajectories and their endpoints (which were verified to be near stable fixed points) are colored by the task on the previous trial and the number of preceding unrewarded trials. While the organization differs between the two networks, both show separation not only by previous and/or to-be-performed task, but also by task certainty, approximated here by number of preceding unrewarded trials. Principal components reflect the dimensions that capture the most variance in the endpoint activity (X markers). **(B)** Activity from stimulus onset (X marker) until the end of a 400 ms inter-stimulus delay period (circle markers) for two randomly selected example trial histories for each network, one after a rewarded trial (top row) and one after 3 or more unrewarded trials (bottom row), where the last chosen task was location. For each example trial history, stimulus response dynamics were simulated for five unique pairs of feature values such that relevant (location, solid black lines) and irrelevant (frequency, dashed black lines) feature axes could be visualized. The location value is indicated by trajectory and marker fill color, and the frequency value is indicated by marker outline color. Principal components were computed from the five trajectories at the beginning of the delay, just following stimulus offset (left, triangle markers) and separately at the end of the delay (right, circle markers). These examples illustrate that for both networks, at the end of the delay (circle markers), trajectories are much less separated by irrelevant feature value (marker outline color) on trials after a reward (top) than after several non-rewards (bottom). This effect is less pronounced at the beginning of the delay (triangle markers), indicating that in these networks it emerges largely from relaxation dynamics (i.e. after the stimulus input shuts off) rather than initial stimulus response (i.e. during the stimulus input). In the correct choice network, feature axes remain orthogonal throughout the delay after unrewarded trials, while in the monkey choice network they become less orthogonal (bottom row).

**Supplementary Figure S8:**
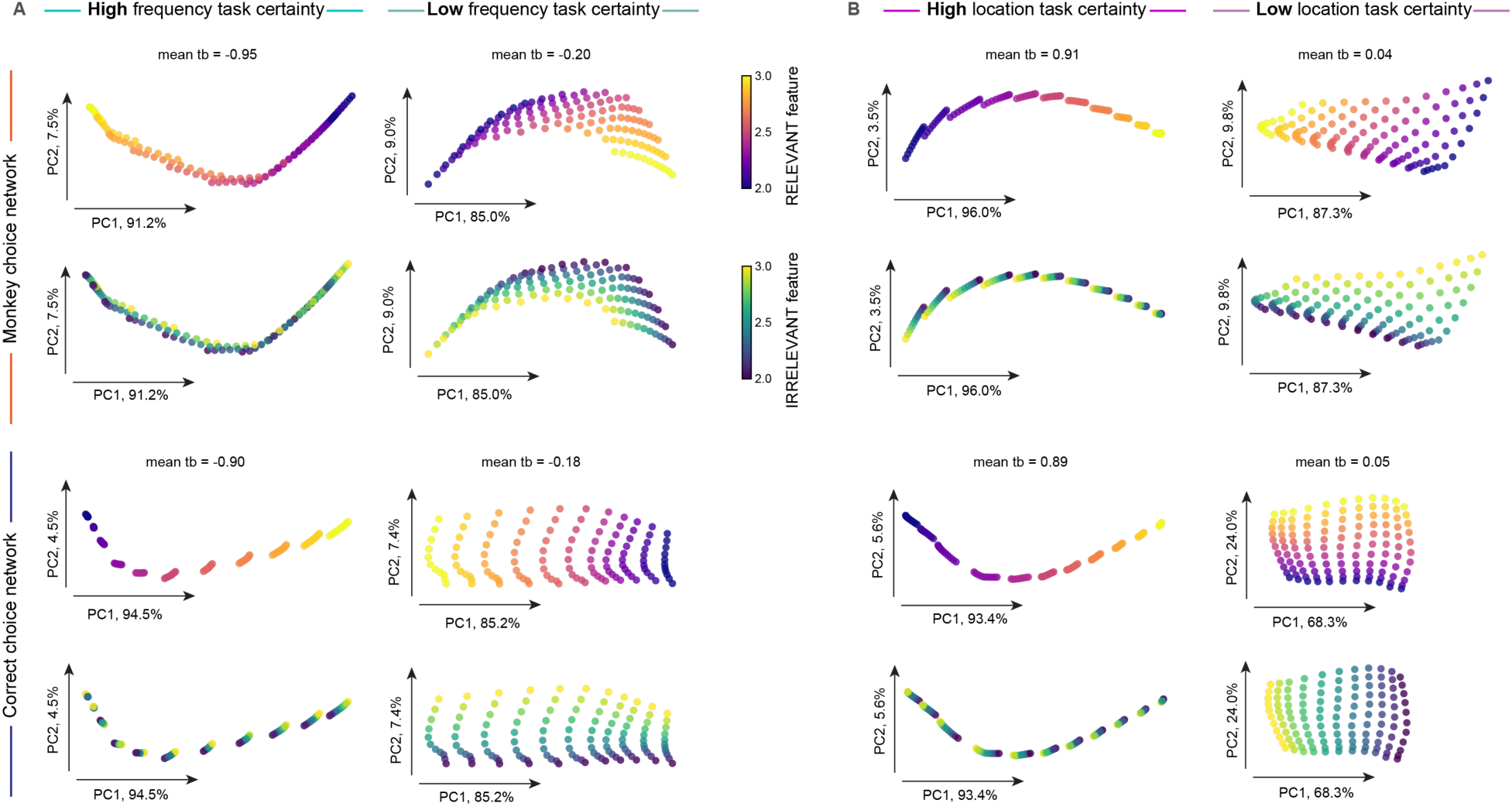
Stimulus maps of RNN hidden layer activity at the end of the inter-stimulus delay for example high and low task certainty conditions, see also Figure 2E. **(A)** Hidden layer activity 100 ms before the onset of the second stimulus during the frequency task. Activation patterns were projected onto the first two principal components (>90% variance), which capture >90% of the variance. Each data point is one of the 11x11 combinations of the initial stimulus. **(B)** Hidden layer activity 100 ms before the onset of the second stimulus during the location task, same convention. In both panels, high task certainty conditions are in the left column, and low task certainty conditions are in the right column. The top rows show the monkey choice model (MCM), and the bottom rows show the correct choice model (CCM). Activity is colored by either the relevant feature value (rows 1 and 3) or the irrelevant feature value (rows 2 and 4). Under high certainty (left columns), the relevant feature is decodable (color gradient along the curve), while the irrelevant feature is not linearly decodable. Under low certainty (right columns), the irrelevant feature becomes more decodable in both models (color is more spread out and separable). However, in the CCM, the relevant and irrelevant feature values vary along orthogonal directions (approximately aligned with the PCs), whereas in the MCM, they interfere. The example trial history inputs were constructed to match the task belief outputs (see S4A) of the two models (labeled “mean tb”). For frequency (A) the trial history inputs were: MCM, low certainty: 7 rewarded trials followed by 2 unrewarded trials where “frequency increase” was chosen and perceptual difficulty was low; CCM, low certainty: same but with 8 rewarded, 1 unrewarded trial (note that this is consistent with the CCM taking less trials to switch tasks (Fig. S3C)); both models, high certainty: same but with all 9 trials rewarded. For location (B) the trial history inputs were: MCM, low certainty: 6 rewarded trials followed by 3 unrewarded trials where “location increase” was chosen and perceptual difficulty was high; CCM, low certainty: same but with 7 rewarded, 2 unrewarded trials; both, high certainty: same but with all 9 trials rewarded.

**Supplementary Figure S9:**
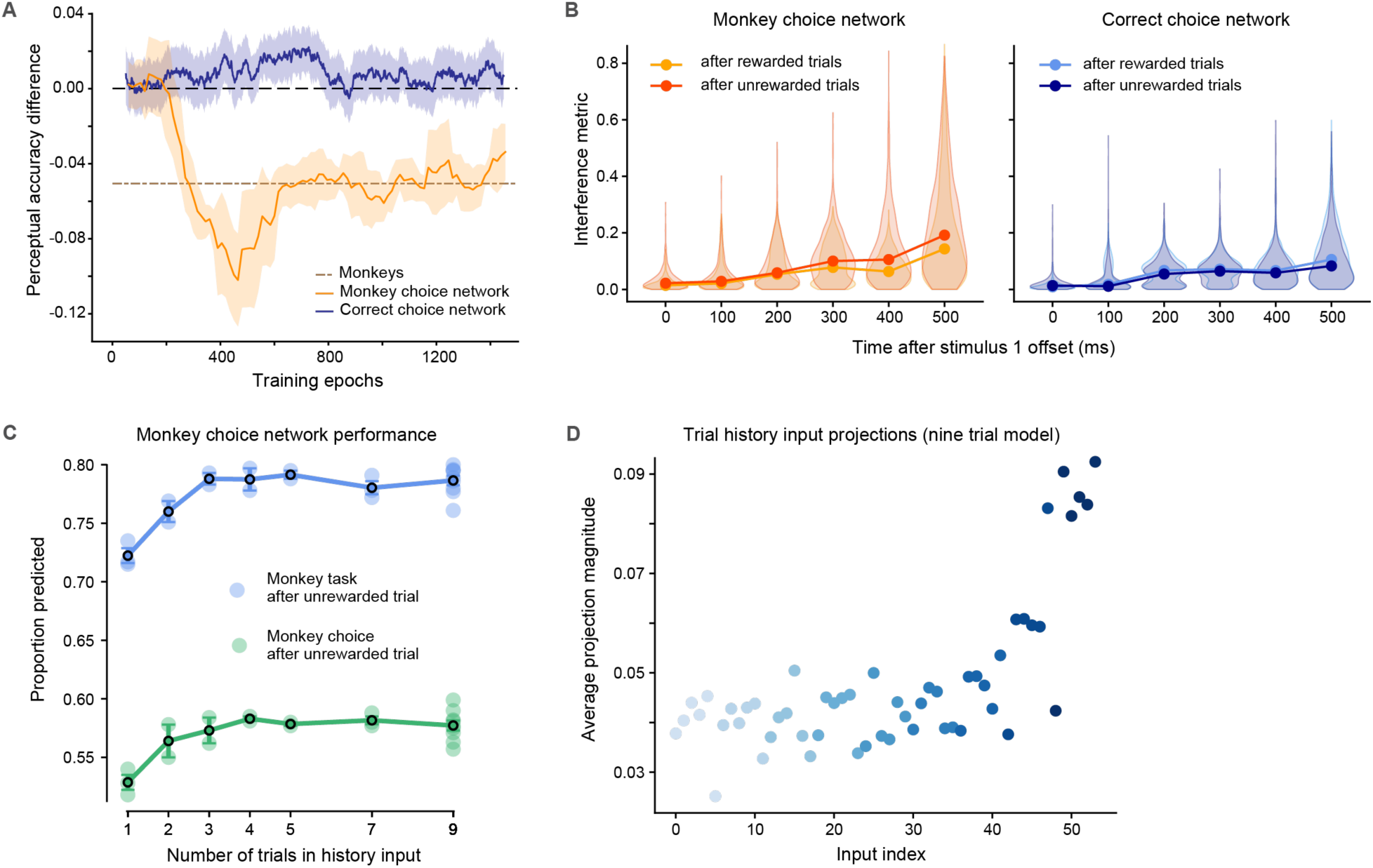
Validation of RNN architecture and trial history window. **(A)** Perceptual accuracy difference (after unrewarded – after rewarded trials) during training for fully connected (non-modular) RNNs. Similar to the modular networks shown in Fig. 2B, the monkey choice network (orange) learns the behavioral drop (monkey average, dashed grey line), while the correct choice network (blue) does not. **(B)** Feature interference metric (DCA) during the inter-stimulus delay for the fully connected networks. As in the modular networks (Fig. 2G), the monkey choice network (left) exhibits significantly higher interference after unrewarded trials (dark orange) than after rewarded trials (light orange). The correct choice network (right) shows low interference in both conditions. **(C)** Predictive performance of the modular monkey choice network as a function of the number of trials used in the history input. The network’s ability to predict the monkey’s choice (green) and task (blue) plateaus after approximately 3-5 trials of history. **(D)** Average projection magnitude of inputs to the task module for the network trained with a 9-trial history window. The network learns to weigh the most recent trials (higher indices, right side) more heavily, as shown by the larger projection magnitudes. Inputs are grouped in sixes (corresponding to perceptual difficulty, one-hot-encoded choice, and reward feedback) for each of the 9 preceding trials.

## Notes

### Competing Interest Statement

The authors have declared no competing interest.

### Summary of Updates

Major manuscript revisions based on reviewer feedback.

